# Phenomenological models of Na_V_1.5. A side by side, procedural, hands-on comparison between Hodgkin-Huxley and kinetic formalisms

**DOI:** 10.1101/720862

**Authors:** Emilio Andreozzi, Ilaria Carannante, Giovanni D’Addio, Mario Cesarelli, Pietro Balbi

**Author notes:** Corresponding Author, address: via Maugeri, 4 - Pavia 27100 ITALY.

## Abstract

**Background:** Computational models of ionic channels represent the building blocks of conductance-based, biologically inspired models of neurons and neural networks. Ionic channels are still widely modelled by means of the formalism developed by the seminal work of Hodgkin and Huxley, although the electrophysiological features of the channels are currently known to be better fitted by means of kinetic (Markov-type) models.

**Objective:** The present study is aimed at showing why kinetic, simplified models are better suited to model ionic channels compared to Hodgkin and Huxley models, and how the manual optimization process is rationally carried out in practice for these two kinds of models.

**Methods:** Previously published experimental data on macroscopic currents of an illustrative ionic channel (Na_V_1.5) are exploited to develop a step by step optimization of the two models in close comparison. The proposed kinetic model is a simplified one, consisting of five states and ten transitions.

**Results:** A conflicting practical limitation is recognized for the Hodgkin and Huxley model, which only supplies one parameter to model two distinct electrophysiological behaviours (namely the steady-state availability and the recovery from inactivation). In addition, a step by step procedure is provided to correctly optimize the kinetic model.

**Conclusion:** Simplified kinetic models are at the moment the best option to closely approximate the known complexity of the ionic channel macroscopic currents. Their optimization is achievable by means of a rationally guided procedure, and it results in models with computational burdens comparable with those from Hodgkin and Huxley models.

## Introduction

Nowadays biologically inspired neural simulations, sustained by both escalating computational power and more detailed comprehension of the nervous system physiology (Churchland and Sejnowski 2016), are becoming increasingly popular and appreciated, despite their concurrent boosting complexity (Hartveit et al. 2018; Arkhipov et al. 2018; Cavarretta et al. 2018; Markram et al. 2015; Kozlov et al. 2014; Traub et al. 2005).

At the core of those simulations, multi-compartmental, conductance-based models of single neural cells with variable degree of morphological and biophysical details can be retrieved. The modelled single cells in turn mainly and directly derive their electrophysiological properties (that is, the fundamentals of the entire modelled neural networks) from the kinetics of the macroscopic currents of different kinds of voltage-gated ionic channels (Hille 1992).

Phenomenological models of ionic channels, therefore, constitute the building blocks of biologically inspired neuronal cells and neural networks.

Current electrophysiological techniques are able to provide huge amount of data with unprecedented details on voltage-gated ionic channels. Developed as patch-clamp methods (Neher and Sakmann 1976), when used in whole-cell configuration, these techniques are most suitable for recording the macroscopic currents of ionic channels, greatly improving our comprehension of the kinetic features of the channels.

Yet, a gap in integrating the improved access to functional properties of ionic channels into whole-cell models has been for long recognized (Cannon and D’Alessandro 2006; Patlak 1991).

Until recently the phenomenological behaviour of the voltage-gated ionic channels has been mainly modelled according to the seminal work of Hodgkin and Huxley (Hodgkin and Huxley 1952a). The availability of analytical solutions for Hodgkin and Huxley (HH) equations, which can be directly fitted to the experimental results, makes the HH formalism a gold standard for ionic channels modelling. In addition, the light computational load of HH models makes them particularly well suited to be implemented in biologically inspired neural networks of increasing complexity. However, the HH formalism turned out to carry theoretical and practical limitations in accurately reproducing the increasing complexity of the electrophysiological behaviour of ionic channels (Bezanilla 2008; Maurice et al. 2004; Meunier and Segev 2002; Strassberg and DeFelice 1993; Patlak 1991).

On the other hand, Markov-type kinetic models, characterized by a set of not independent states with transitions between states governed by barrier-style equations, proved capable to approximate with higher accuracy the known complexity of the voltage-gated ionic channels (Destexhe and Huguenard 2010; Borg-Graham 1999; Destexhe et al. 1994; Kuo and Bean 1994). Markov-type kinetic models can also be designed to reproduce the behavior of a single channel protein, with each state corresponding to a specific physical conformation of the protein. These extremely detailed models, however, needs a huge number of states (e.g., Börjesson and Elinder 2008), resulting in a heavy computational load, which makes them unsuitable for implementation in neural networks.

Thus, simplified kinetic models have been proposed (Destexhe and Huguenard 2010; Destexhe et al. 1994) to provide efficient modelling of ion-channels behaviour, by overcoming the limitations of the HH formalism, while keeping the computational burden sufficiently low to make them suitable for implementation in biologically inspired neural networks.

They are built with a reduced number of states and addressed to deterministically reproduce the macroscopic current kinetics of ionic channels, rather than their conformational changes.

In this work, a point-by-point comparison between HH and kinetic models of an illustrative human sodium channel (Na_V_1.5) is performed, with the aim of clarifying the process of optimization of a simplified kinetic model in close relationship with a correspondent HH model. The study unveils the limits of HH models and suggests simplified kinetic models as an essential tool to *in silico* approximate the known complexity of the ion channels kinetics. A critical limitation of HH formalism resulted from the simultaneous optimization of the steady-state availability and the recovery from fast inactivation, which led to a flawed modelling of the electrophysiological features of the channel.

## Methods Experimental data of Na_V_1.5 (Zhang et al. 2013)

We chose to simulate the electrophysiological behaviour of Na_V_1.5 because quite comprehensive experimental data for this channel have been already published and made available to be shared and reused under a Creative Common license (Zhang et al. 2013).

Experimental data on Na_V_1.5 macroscopic currents were obtained by heterologously expressing the α-subunit of the ionic channel in a mammalian cell line (Human Embrionic Kidney 293 cells). No β-subunits were co-expressed in the study and the electrophysiological experiments were conducted by means of the whole-cell patch-clamp method at room temperature (Zhang et al. 2013).

Na_V_1.5 is the isoform of the sodium channel α-subunit typically expressed in the heart, where it is mainly involved in the cardiac rythmogenesis, as revealed by the rare channel mutations responsible of severe genetic arrythmiae (Southan et al. 2016; Zhang et al. 2013). But Na_V_1.5 were also detected in different structures of the brain (Wu et al. 2002), where they clustered at a high density in the neuronal processes, mainly axons.

### The HH model

According to the original formulation of Hodgkin and Huxley (Hodgkin and Huxley 1952a), the HH model is based on a membrane equation describing three ionic currents in an isopotential compartment:

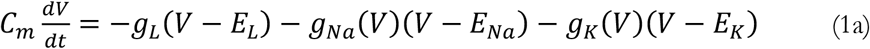

where *C_m_* is the membrane capacitance, *V* is the membrane potential, *g_L_*, *g_Na_* and *g_K_* are the membrane conductances for leak currents, *Na*^+^ and *K*^+^ currents respectively, *E_L_*, *E_Na_* and *E_K_* are their respective reversal potentials.

Hodgkin and Huxley hypothesized that ionic currents result from the assembly of several independent gating (or carrier) particles which must occupy a given position in the membrane to allow the ions to flow (Hodgkin and Huxley 1952a). Each gating particle can be on either side of the membrane and bears a net electronic charge such that the membrane potential can switch its position from the inside to the outside or vice-versa.

The transition from these two positions is therefore voltage-dependent, according to the diagram:

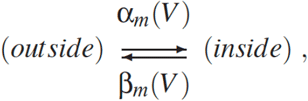

where *α* and *β* are respectively the forward and backward rate constants for the transition from the outside to the inside position in the membrane. If *m* is defined as the fraction of particle in the inside position, and (1−*m*) as the fraction outside the membrane, the first-order kinetic equation can be obtained:

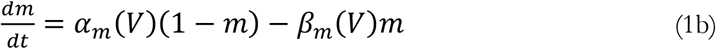

Assuming that particles must occupy the inside position to conduct ions, then the conductance must be proportional to some function of *m*. In the case of *Na*^+^ current in the squid giant axon, Hodgkin and Huxley (1952a) found that its nonlinear behavior, its delayed activation and the sigmoidal rising phase were best fit by assuming the conductance proportional to the product of two variables (Hodgkin and Huxley 1952a):

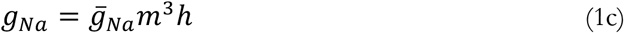

where *ḡ_Na_* is the maximal value of the conductance, and *m* and *h* represent the fraction of two different types of gating particles. The interpretation is that the assembly of three gating particles of type *m* and one of type *h* is required for *Na*^+^ ions to flow through the membrane. These particles operate independently of each other, leading to the *m^3^h* form.

Long after the work of Hodgkin and Huxley, it was established that ionic currents are brought about by the opening and closing of ion channels, and the carrier particles were reinterpreted as gates inside the pore of the channel. Thus, the reinterpretation of Hodgkin and Huxley’s hypothesis was that the pore of the channel is controlled by four internal gates, opening independently of each other, and that all four gates must be open in order for the channel to conduct ions.

The rate constant *α*(*V*) and *β*(*V*) of *m* are such that depolarization promotes opening the gate, a process called *activation*. On the other hand, the rate constants of *h* are such that depolarization promotes closing the gate (and therefore closing of the entire channel because all gates must be open for the channel to conduct ions), a process called *inactivation*.

Thus, the set of differential equations (HH equations) able to explain the *Na*^+^ current characteristics is made by the (1b) and the following one:

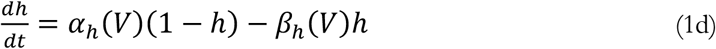

The rate constants *α_m_, β_m_, α_h_* and *β_h_* were estimated by fitting empirical functions of voltage to the experimental data (Hodgkin and Huxley, 1952a):

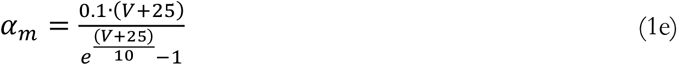

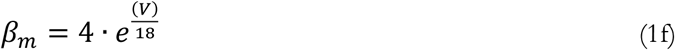

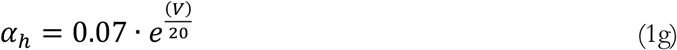

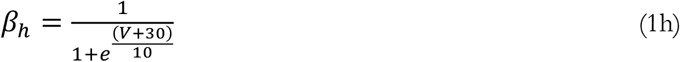

These functions were estimated at a temperature of 6°C and the voltage axis was reversed in polarity (voltage values were given with respect to the resting membrane potential).

By adopting the modern convention on voltage axis orientation and a resting voltage of −65 mV, the HH equations can be redrawn:

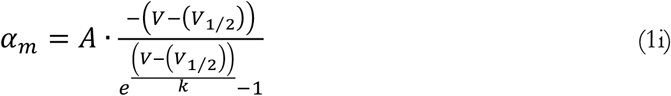

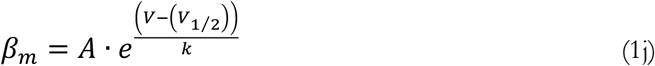

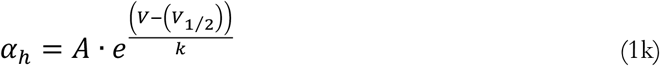

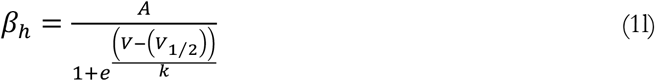

with the following parameters of the forward and backward rate constants able to replicate the original Hodgkin and Huxley (1952a) experimental values (Fig 1A-B):

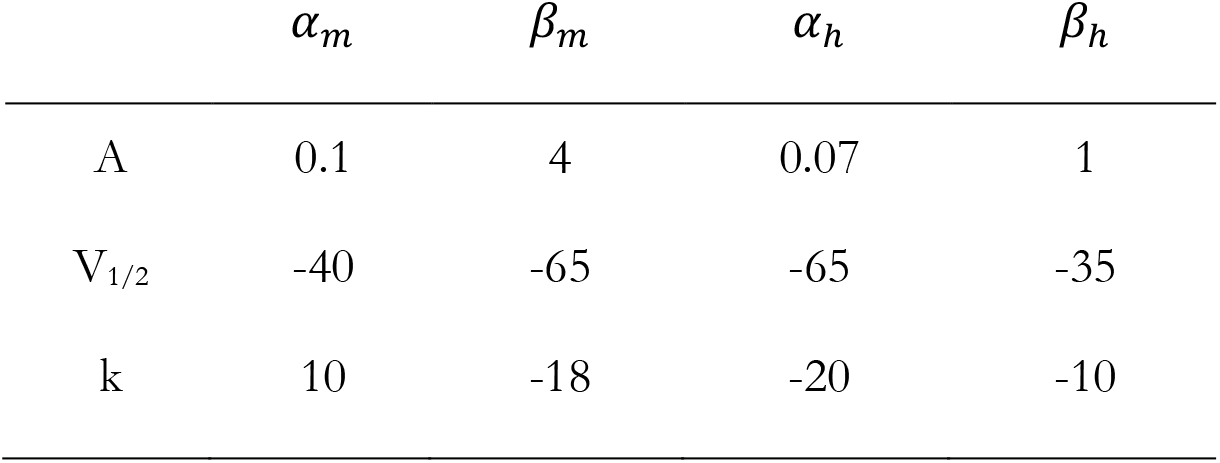

| | $\alpha_m$ | $\beta_m$ | $\alpha_h$ | $\beta_h$ |
| --- | --- | --- | --- | --- |
| A | 0.1 | 4 | 0.07 | 1 |
| $V_{1/2}$ | -40 | -65 | -65 | -35 |
| k | 10 | -18 | -20 | -10 |

**Figure 1.**
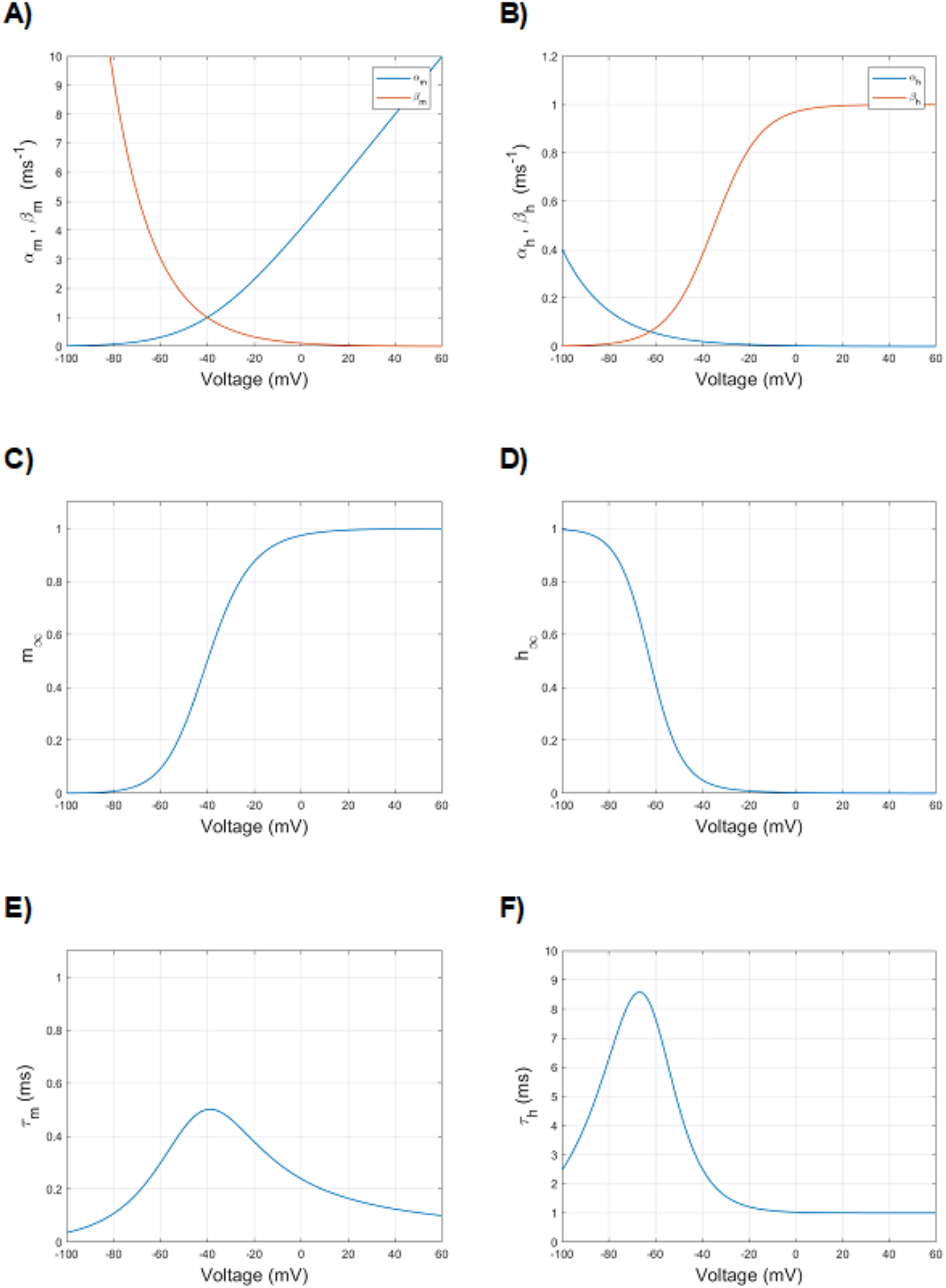
Voltage dependence of particle (gate) variables and rate constants of sodium ionic channel of *Loligo*, as found in the original paper by Hodgkin and Huxley (Hodgkin and Huxley, 1952a), redrawn according to the actual notation on voltage axis direction and rest potential reference. A) Voltage dependence of forward (*α_m_*, blue) and backward (*β_m_*, red) rate constants of activation. B) Voltage dependence of forward (*α_h_*, blue) and backward (*β_h_*, red) rate constants of inactivation. C) Voltage dependence of steady-state activation (*m*_∞_). D) Voltage dependence of steady-state inactivation (*h*_∞_). E) Voltage dependence of activation time constant (*τ_m_*). F) Voltage dependence of inactivation time constant (*τ_h_*).

The HH model is often written in an equivalent form, more convenient to fit the experimental data:

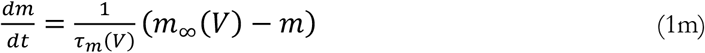

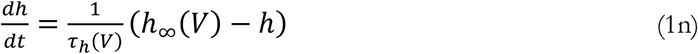

where

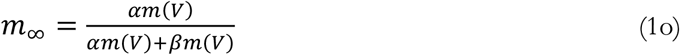

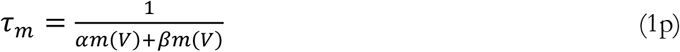

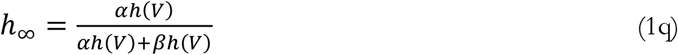

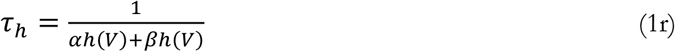

Here, *m*_∞_ is the steady-state activation and *τ_m_* is the activation time constant of *Na*^+^ current (Fig 1C and Fig 1E). In the case of *h, h*_∞_ and *τ_h_* are steady-state inactivation and inactivation time constant, respectively (Fig 1D and Fig 1F).

### The kinetic model

The kinetic model, derived by a previously developed one (Balbi et al. 2017), has a five-state diagram, with one open, two closed and two inactivated states (Fig 2).

**Figure 2.**
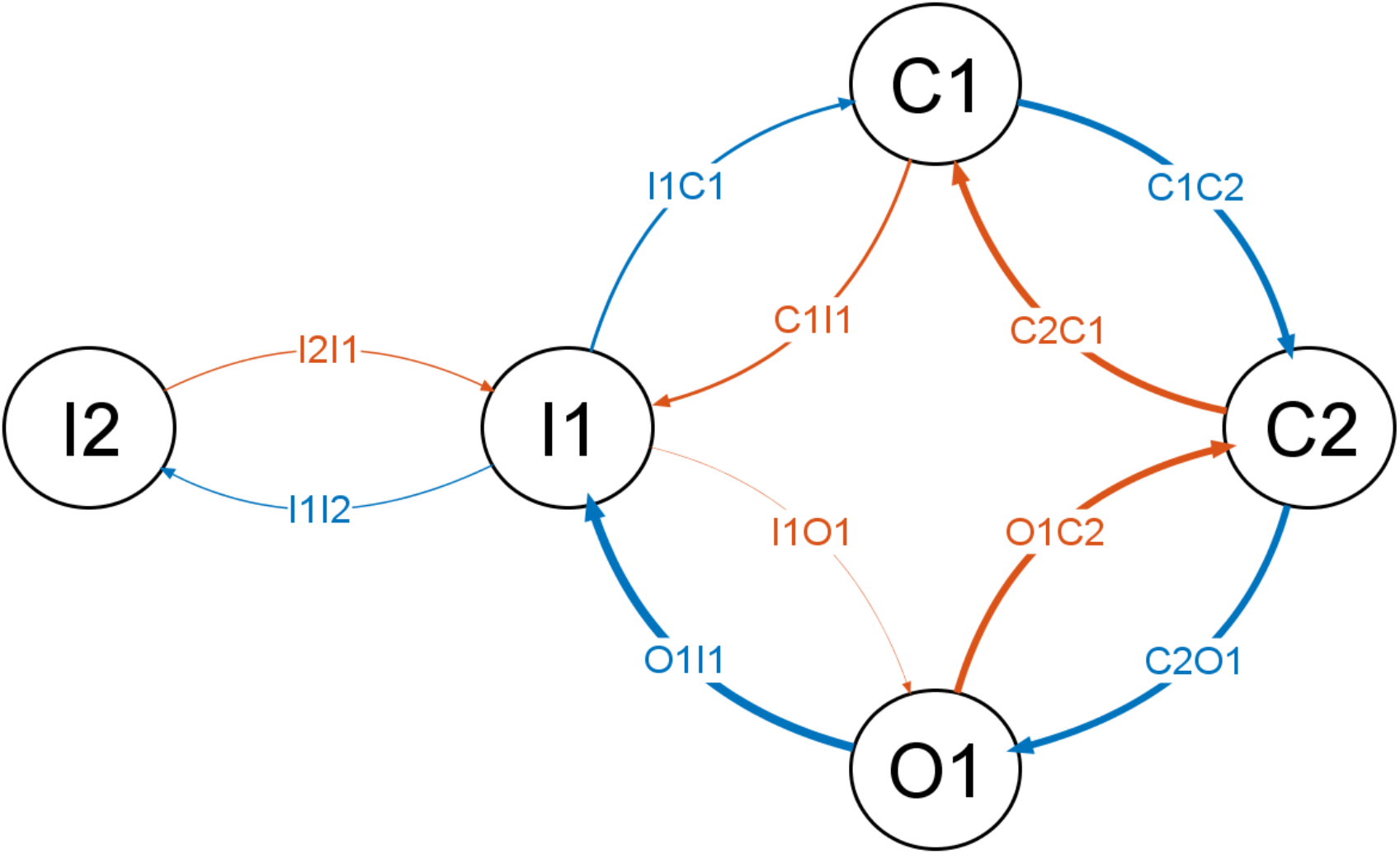
Diagram of the five-state kinetic model. The thickness of the transitions is drawn as a schematic cue to the maximal amplitude of the transition rate.

The second inactivated state (I2) is considered as a deeper inactivated state than I1, only connected to I1.

All transitions between two consecutive states are considered reversible, with one exception (see below), and the paired forward and backward transitions were computed by equations carrying numerical values (coefficients) of the same order of magnitude.

The only exception was the O1 to I1 transition, which could be considered irreversible since its backward transition (I1 to O1) was characterized by an extremely small rate as compared to the other transitions’ ones.

The dynamics of the fractions of channels being in the five different states (later referred to as the “states”, for the sake of simplicity) are described by the following set of coupled ordinary differential equations:

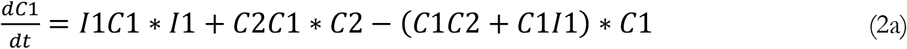

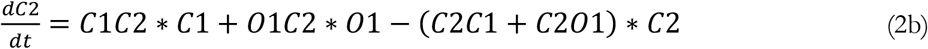

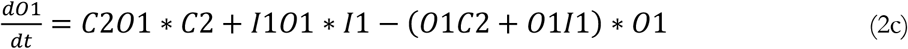

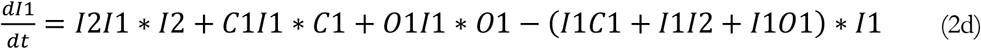

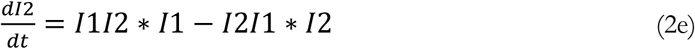

Moreover, the states obey the law of mass conservation:

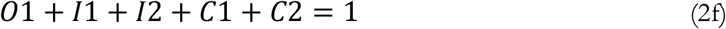

The ionic channel current is governed by Ohm’s law, wherein channel conductance is proportional to the fraction of channels in the open states with the sodium maximal conductance being the proportionality coefficient, according to the following equation:

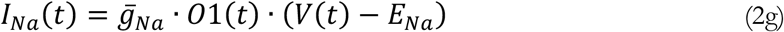

where *ḡ_Na_* is the sodium maximal conductance, O1 is the fraction of channels in the open states (bounded between 0 and 1) and *E_Na_* is the reversal potential of the sodium ion.

Since the studies by Hodgkin and Huxley (Hodgkin and Huxley 1952a), the voltage dependence of the rate transitions has been mathematically modelled as an exponential function, or as a sigmoid, or as a combined linear and exponential function (Figg 1A-B). In other cases, according to the theoretical approach of the thermodynamic theory (Destexhe et al. 1994; Borg-Graham 1999), a sigmoid curve with minimum and maximum asymptotes has been adopted. It was described by the following equation

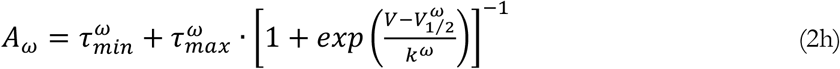

where *ω* is the transition between two states, 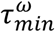 and 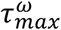 are the two asymptotes, 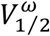 the hemiactivation voltage, and *k^ω^* the slope which describes the voltage sensitivity of the transition rate.

In a previous work (Balbi et al. 2017) we found that the most suitable general equation to be adopted in all the transitions was a sigmoidal one. For most of the transitions, the minimum asymptote was conveniently set to zero, while in few cases, notably for the O1 to I1 transition, it needed a non-zero value. Furthermore, to accurately accommodate the time course of the current-voltage curves, it was found appropriate to slightly modify the sigmoid, adding a bending at the beginning of the rising slope of the curve (Figg 3A-I).

**Figure 3.**
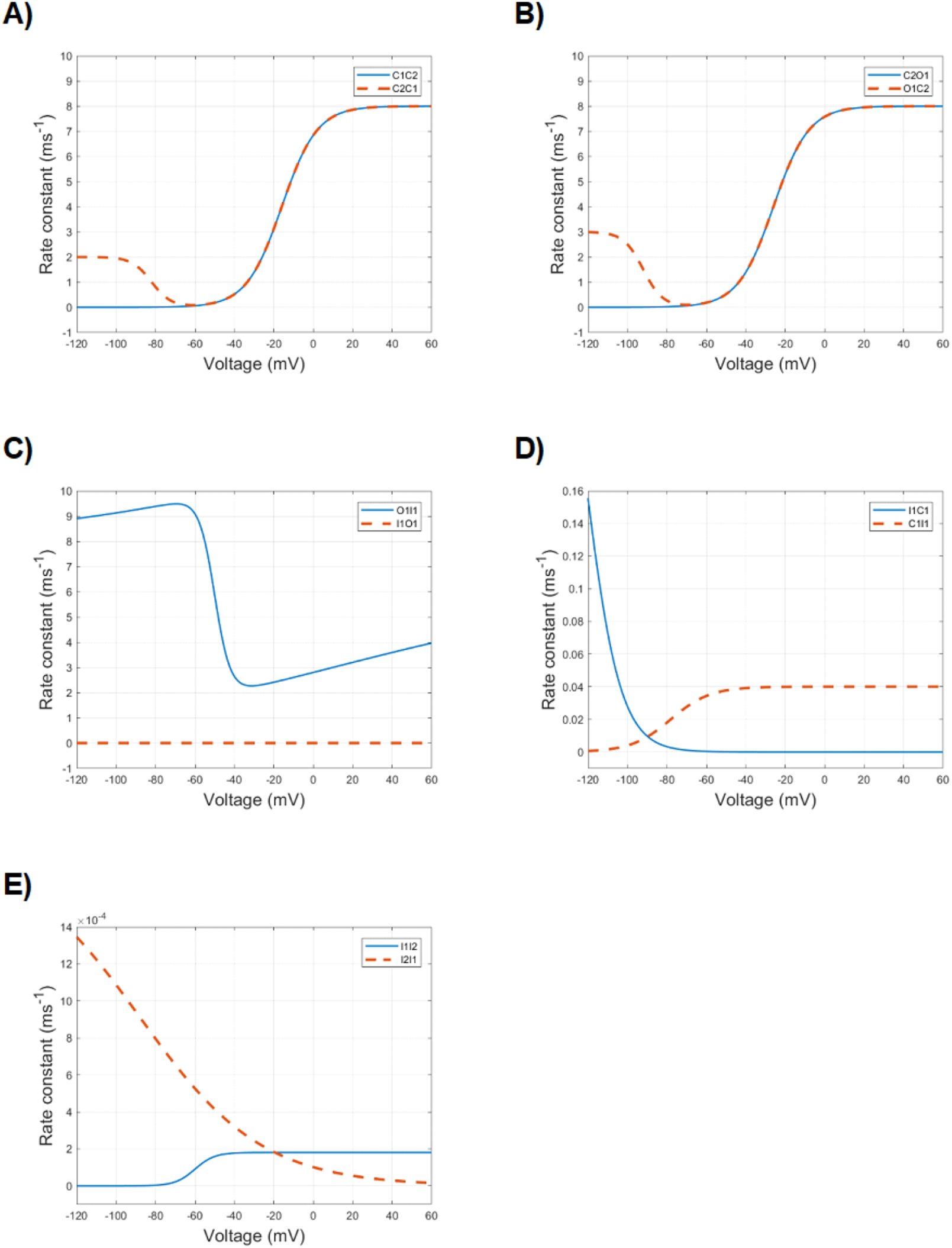
Voltage dependence of the following ten transition rates of the kinetic model: A) C1C2 (blue) and C2C1 (red); B) C2O1 (blue) and O1C2 (red); C) O1I1 (blue) and I1O1 (red); D) I1C1 (blue) and C1I1 (red); E) I1I2 (blue) and I2I1 (red). Note the different ordinate scales.

In this way, the modified sigmoid could be mathematically described as the combination of two sigmoids with opposite slope: the first one describing the transition rate for more polarized voltages and carrying a positive slope factor, the second one providing the transition rate values for more depolarized voltages and carrying a negative slope factor. As a result, the general equation adopted to describe this double sigmoid was set as

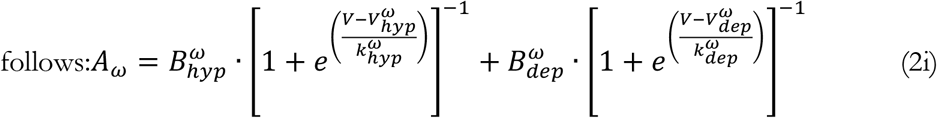

where 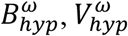, and 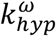 are, respectively, the magnitude, the hemiactivation and the slope factor, of the voltage dependence of the transition rate *ω* in the hyperpolarized region, and 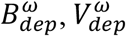 and 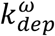 are the corresponding values in the depolarized region. With this formalism, the slope factor (*k*) assumes a positive value in the hyperpolarized region and a negative value in the depolarized one. In addition, when the transition rate is better described by a simple sigmoid, which is the case in most of the transitions, one of the two terms of the general equation can be conveniently dropped.

### The electrophysiological protocols

#### Activation curves and normalized conductance-voltage dependence

Voltage-clamp intensity-voltage (activation) curves are obtained by sequentially voltage clamping the channel in steps of 5 or 10 mV from a resting value (Fig 4A).

**Figure 4.**
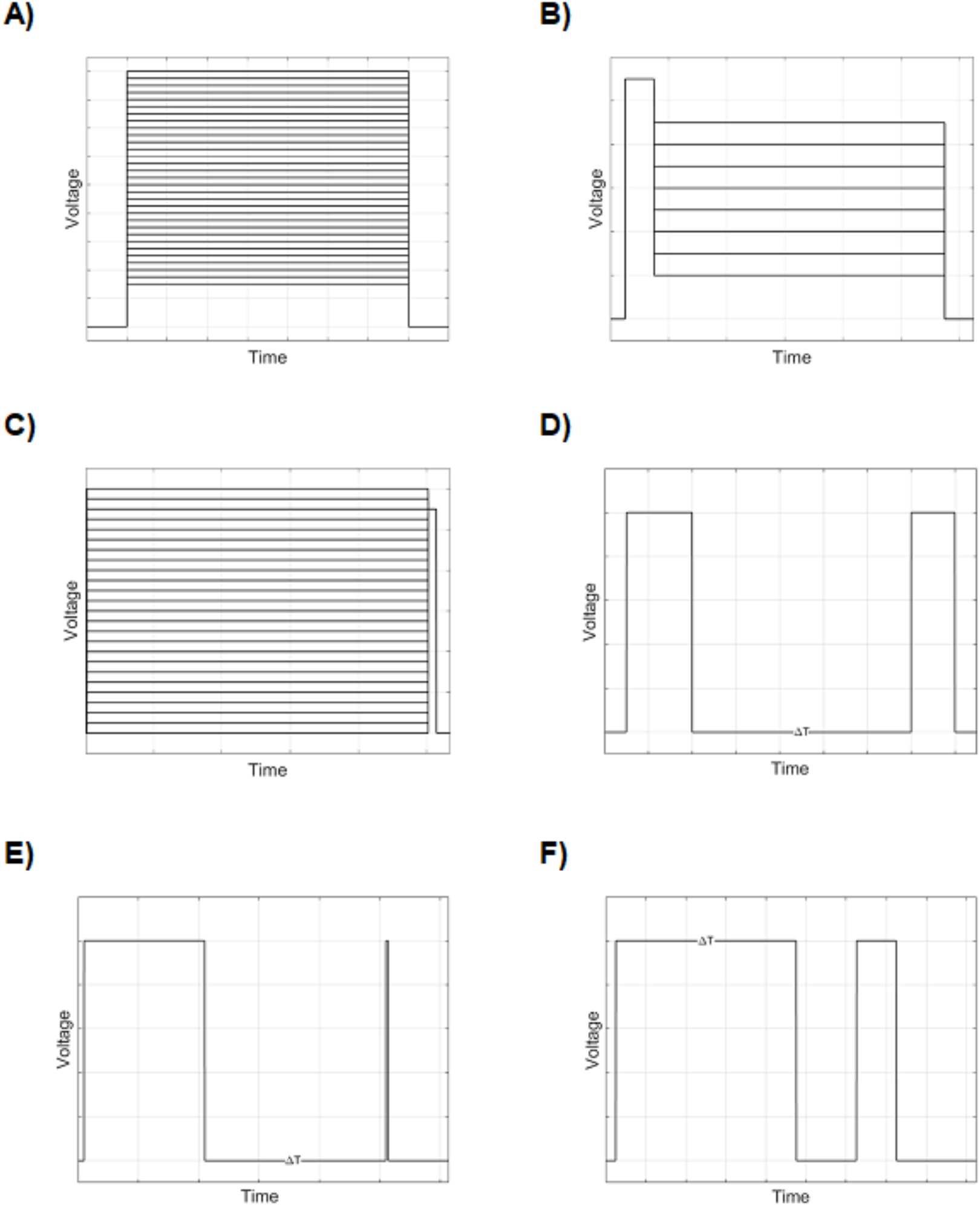
Experimental and simulated voltage clamp protocols. A) Activation: A 2-ms pulse at −120 mV is followed by a series of 14 ms long depolarizations (from −90 to +60 mV), in steps of 5 mV. B) Deactivation: After a 0.5-ms pulse at −120 mV, a 0.5-ms depolarization at −10 mV is delivered, followed by a series of 5-ms long repolarization, from −100 to −30 mV, in steps of 10 mV. C) Steady-state availability: Following a series of 500-ms long conditioning depolarizations from −120 to 0 mV in steps of 5 mV, a 20-ms long test stimulus at −10 mV is delivered. D) Recovery from fast inactivation: After a 30-ms long conditioning depolarization at −20 mV (P1), variable time intervals (from 0.1 to 1000 ms) of repolarization at −120 mV are followed by a 20-ms long probing depolarization at −20 mV (P2). E) Recovery from slow inactivation: similar to the previous one protocol, with the difference that the conditioning pulse (P1) is 1000 ms long, and the repolarization time intervals are from 0.1 ms to 7000 ms. F) Development (onset) of slow inactivation: A series of depolarizations at −20 mV of increasing duration, from 10 to 10000 ms (P1), is followed by a brief (30 ms) repolarization impulse at −120 mV and then by a 20-ms long test depolarization at −20 mV (P2).

Normalized conductance-voltage relationship is obtained by converting the current peak values into the respective conductance values, according to the equation (3a),

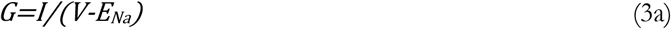

where *G* is the conductance, *I is* the peak current, *V* is the membrane voltage, *E_Na_* is the sodium equilibrium potential, and *V-Ema* is referred to as the driving force. The so evaluated conductance is plotted against voltage clamp values, and the conductance-voltage curves (Fig 6A) are usually fitted to a Boltzmann equation (3b):

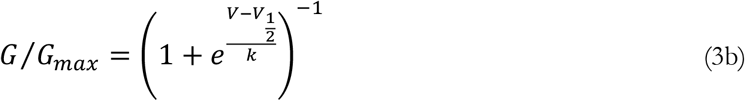

where *G_max_* is the maximum conductance, *V_1/2_* is the half maximal voltage and *k* is the slope factor.

For each single activation curve, the time constants of activation (from the onset of the current to the peak) and inactivation (decay from activation, from the peak to the steady-state) are calculated. According to the work by Hodgkin and Huxley (1952a), the time constants of activation and inactivation are derived by fitting the entire simulated curve to the following equation:

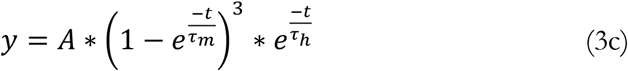

where *τ_m_* and *τ_h_* are the time constant of activation and inactivation, respectively. In this case, the activation segment of the curve is fitted by a third power exponential, while the inactivation (decay) one by a simple exponential.

#### Deactivation curves

The so-called tail currents are evoked by a quick repolarization after a brief depolarizing pulse (Fig 4B). The preliminary brief depolarizing pulse causes a (maximal) fraction of channels to open, and the following depolarization is administered before inactivation is fully deployed. Thus, the protocol is designed to sample the return of the fraction of *m* gating particles to low values during repolarization, in terms of HH formalism, or, in terms of kinetic model, to sample the transition between the open and closed states, a process named deactivation. It is worth noting that deactivation differs from inactivation, as the latter defines the transition between open and inactivated states.

By varying the voltage of the repolarizing pulse, a series of curves of tail currents are obtained which can be fitted to the mono-exponential equation (3d)

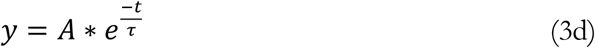

#### Voltage dependence of (normalized) current during fast inactivation (steady-state availability)

In steady-state availability protocols (Fig 4C), conditioning pulses of variable voltage and long duration are administered to reach a steady-state condition, before applying a depolarizing test stimulus of fixed amplitude. The conditioning stimulus is able to set a fraction of channels into a steady inactivated state, and the following depolarizing test stimulus samples the fraction of channels available to open. It is worth mentioning that the inactivation occurs even before the activation threshold is reached. In other words, depolarizing stimuli below the activation threshold are able to move a fraction of channels from closed to inactivated states, without passing through the open state. The normalized current peaks following the test stimuli are plotted against the voltage of the conditioning stimuli, and the resulting curve is fitted to the Boltzmann equation (3e)

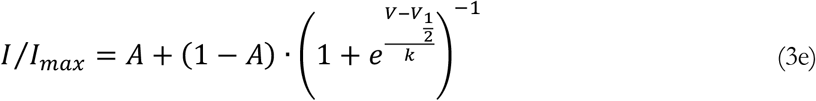

where *I* is the peak current, *I_max_* is the maximal peak current, *A* is the fraction of noninactivating channels, *V_1/2_* is the voltage at which half of the channels are inactivated, and *k* is the slope factor.

#### Recovery from fast inactivation (repriming)

The recovery from inactivation (or repriming) is sampled by a depolarizing (conditioning) pulse (P1) followed by a variable time interval (from tenths to hundreds of milliseconds, in the case of Na_V_1.5; Zhang et al. 2013) of repolarization, and then by a second depolarizing pulse (P2), which tests the fraction of channels that recovered from the inactivation (Fig 4D). A normalized intensity-time curve is obtained by plotting the current peak of the test stimuli (compared to the pre-conditioning stimulus: P2/P1) against the duration of the repolarization. The derived curve can be fitted to a single exponential function:

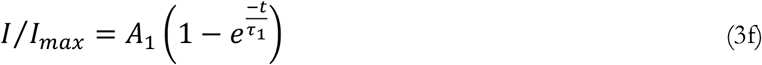

#### Development of slow inactivation

An initial depolarizing conditioning pulse (P1, −20 mV) of increasing duration (Δt, from 10 ms to 10 s) is followed by a brief (30 ms) repolarizing pulse and then by a second depolarizing test impulse (P2), to probe the fraction of non-inactivated, available channels (Fig 4E).

For short durations, the conditioning pulse initially is able to put a fraction of the channels into the (fast) inactivation. This inactivation features fast kinetics and, in fact, the short duration of the repolarizing pulse is long enough for the channels to recover into non-inactivated states; the second test pulse, then, evokes a full amplitude current. However, for increasing durations of the conditioning pulse P1, the test pulse P2 evokes progressively smaller currents. This phenomenon is commonly interpreted by admitting that progressively longer conditioning tests are able to move a fraction of the channels into a deeper inactivated state with slower kinetics (both for entry and recovery), thus making these channels not available to be opened by the probing test pulse.

The obtained curve is fitted to a single exponential equation (14):

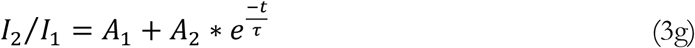

#### Recovery from slow inactivation

Similarly to the fast inactivation, also the recovery from slow inactivation is sampled by means of a double pulse protocol (Fig 4D), in which the first depolarizing conditioning pulse (P1) is followed by a repolarizing time interval of increasing duration (Δt, from 0.1 s to 10 s), followed by a short probing depolarization (P2). The difference with the fast inactivation protocol is that P1 is much longer (1000 ms, compared to 100 ms).

The obtained curve is fitted to a double exponential function (15):

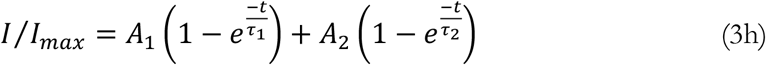

where *A_1_* and *A_2_* are proportional coefficients, *t* is the time, *τ_1_* and *τ_2_* are the fast and slow recovery time constants, respectively.

### Simulations of the experimental procedure in NEURON

All simulated experiments were performed by means of NEURON version 7.6 simulation environment (Carnevale and Hines 2006). The equations of the channel models were written and solved directly using the NMODL language of NEURON, which is a derivative of the MODL description language of the SCoP package (Kohn 1989).

It is worth observing that in NMODL different setups and methods can be used to solve the models. For example, in the DERIVATIVE block the differential equations are actually specified while in the KINETIC block they don’t need to be written, but just the chemical reactions are specified. This is very convenient when the model contains several reactions, although it slows down the simulations. In the present study we preferred to use the same block (DERIVATIVE block) for both models, and the same numerical method to solve them, in order to achieve a reliable comparison.

All virtual experiments were performed on a one-compartmental cylindrical ‘soma’ 50 μm long with a diameter of 63.66 μm, so that the membrane area was set to 10’000 μm^2^. The membrane capacitance was set to 1 μF/cm^2^ (Gentet et al. 2000). The maximal conductance density for each voltage-gated sodium channel isomer inserted into the soma was arbitrarily set to 0.1 S/cm^2^, and the resulting ionic current density was measured in mA/cm^2^. The capacitive currents were subtracted from the total current in all the simulations. The time for single integration step (dt) was set to 0.025 ms for all simulations, but for the running time tests, where the dt was set to 0.001.

The thermal sensitivity of any biological process can be described by its temperature coefficient (Q10). Classically, it is defined as the ratio of a reaction rate measured at two temperatures 10 degrees apart (Bělehrádek 1935). In ion channel research, instead of reaction rates, current amplitudes or time constants are often used to calculate Q10 value. A single value of Q10 is typically reported to indicate the temperature dependence of a channel. In the present study, at every step the rate constants of each transition were multiplied by the temperature coefficient, Q10, calculated as follows:

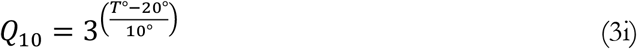

Original NEURON source code was developed to simulate the protocols needed to yield the electrophysiological features of the channels.

The source code along with the virtual experimental procedures is available as a ModelDB (McDougal et al. 2017) entry (access number: 257747).

The simulations were performed on an iMac desktop computer running a MacOS version 10.14.3 (™ and © 1983-2017, Apple Inc, Cupertino, CA, USA) and on a Tuxedo laptop with processor Intel^®^ Core™ i7-7700HQ and 16GB of RAM running Ubuntu 18.04.2 LTS.

### Fitting the experimental data with the models

For modellers, the process of fitting the experimental data can sensibly differ according to the availability of raw electrophysiological data (the single electrophysiological curves of different protocols).

When raw data are available, for the HH models it is possible to apply the procedure devised by Hodgkin and Huxley (Hodgkin and Huxley 1952a). In this case, the *m*_∞_, *τ_m_*, *h*_∞_ and *τ_h_* variables can be directly derived from the single curves of activation, with some assumptions, and the rate constants α_*m*_, β_*m*_, α_*h*_ and β_*h*_ can be calculated by rearranging the equations (1o) to (1r). Afterwards, each single rate constant can be plotted against the voltage and fitted to the original HH expressions (1i) to (1l). Finally, a direct superimposition of real and simulated traces provides the best evidence of the good fit of the model to the experimental data.

However, modellers are rarely provided with raw electrophysiological data, so in most cases it is not possible to directly derive the *m*_∞_, *τ_m_*, *h*_∞_ and *τ_h_* variables. In these cases, channels modelling relies on indirect data about the kinetics of the channels, such as the normalized conductance-voltage relationship, the steady-state availability curve, the repriming curve, etc. By empirically (manually) tuning the parameters of the expressions (1i) to (1l), the best approximation is searched for, in order to reproduce the parameters of the equations (3b) to (3h), which carry substantial information on the macroscopic current kinetics of the channel.

The same procedure is also usually performed for Markov-type kinetic models and has been performed in the present study as well.

### Fitting procedure in NEURON

The developed code automatically supplied the appropriate graphics, which replicated the macroscopic currents and the electrophysiological relationships found in the experimental studies. A both empirical and quantitative curve fitting method was then adopted to reconcile experimental and modelled data. Firstly, the curves and relationships obtained by the simulations were compared by visual inspection to the experimental ones. Then, the modelled curves were fitted to the equations (3b) to (3h), as appropriate, by using a nonlinear least-squares minimization method available in NEURON (Multiple Run Fitter subroutine), which, in turn, derives from the PRAXIS (principal axis) method described by Brent (Brent 1976). Finally, the parameters of the equations (3b) to (3h) of the modelled curves were compared to the experimental ones (Table 1). The agreement of the modelled data with the experimental ones was considered acceptable when the formers were within two standard deviations of the latter.

**Table 1.**
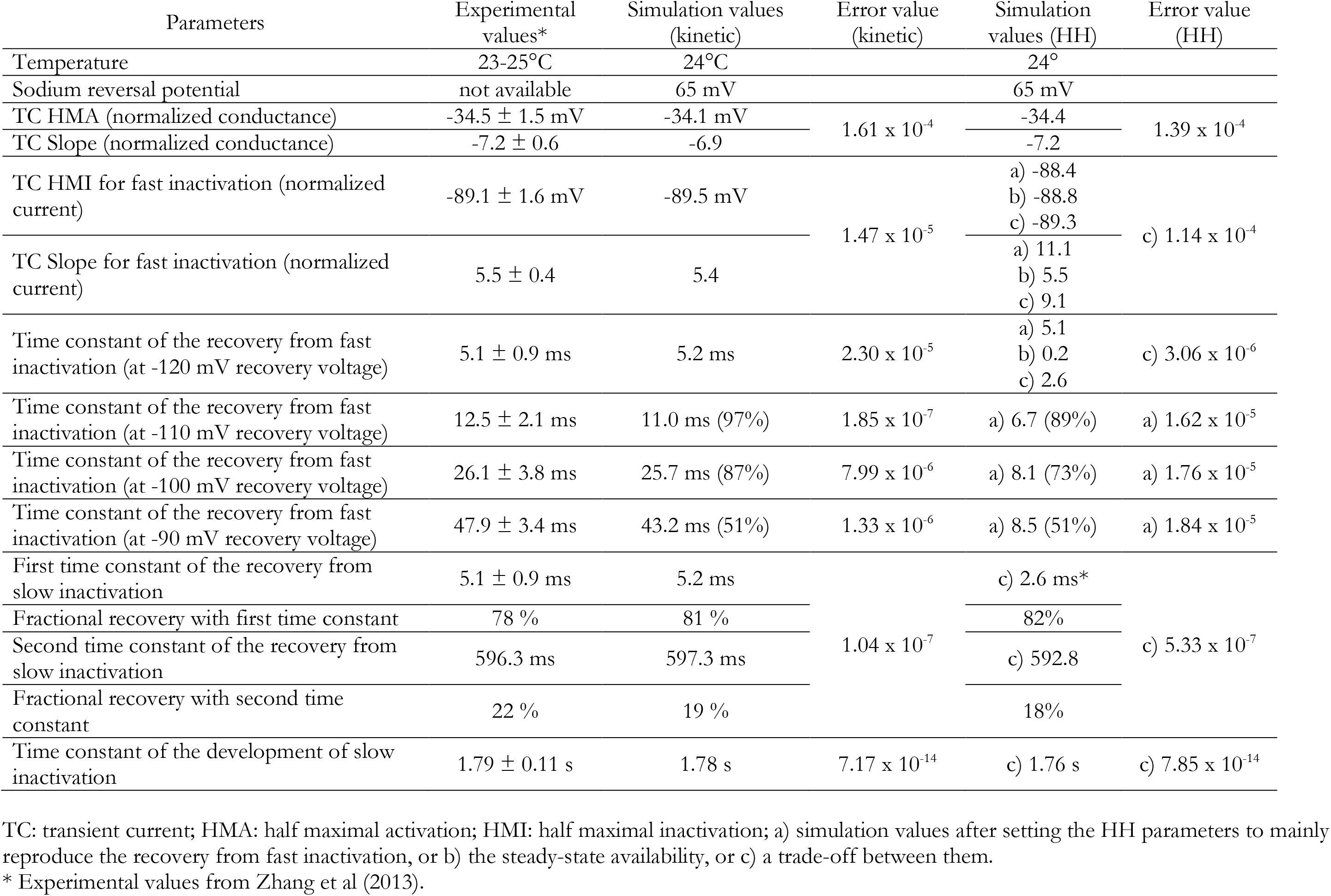
Experimental and simulated values for different protocols performed with HH and kinetic channel models

### Implementation of the channel models in a neuron model

In order to test the suitability of both the kinetic and the HH models to be implemented in cell models, to compare the features of the spikes they provides, and to detect differences in running time and computational load, we inserted the developed channel models in a previously published cell model (Dodge and Cooley 1973) and performed a series of simulations with voltage-clamp and current stimulation.

The previously published cell model was downloaded from the ModelDB (McDougal et al. 2017) repository (accessed on February 15^th^ 2019), where it is accessible with the accession number: 3805.

## Results

Direct comparison of all electrophysiological features of both HH and kinetic model with the experimental data (by Zhang et al. 2013) is provided in Table 1. The displayed parameters supply the best fit to the real data for both models.

The sequential procedures adopted to achieve the best fitting parameters are described in the following paragraphs.

### Activation

#### HH model

At the resting polarized potential, the rate constants of inactivation *α_h_* and *β_h_* (Fig 1B) make the value of *h* close to 1 (that is, no inactivation: Fig 1D). However, the channel is not conducting, as the rate constants of activation make the value of *m* equal to 0 (Fig 1A and Fig 1C). By stepping the potential to more depolarized values, *m* quickly reaches non-zero values and the channel starts conducting ions. At the same time, due to the predominance of backward rate constant of inactivation at depolarized values (Fig 1B), *h* begins to decrease till to a value equal to 0 (inactivation sets in, Fig 1D). However, due to the slower development of inactivation compared to the activation (smaller values of rate constants of *h* compared to those of *m*), inactivation sets in with a delay, which is responsible of the peak of sodium current during activation curves.

The HH original equations for sodium channel (Table 2a, Fig 5B), when used in simulations at 23-24°C (the usual experimental room temperature) and corrected by a Q10 temperature coefficient equal to 3 (Hodgkin and Huxley 1952a), brought extremely fast and unrealistic activation and inactivation time constants for human sodium channels (Fig 5C). By reducing the amplitude of the rate constants of *m* and *h*, as well as by slightly modifying the voltages and slopes parameters of the inactivation rate constants (Table 2b), it is possible to reach an overall initially acceptable fitting for the intensity-voltage curves of Na_V_1.5 (Fig 5D).

**Figure 5.**
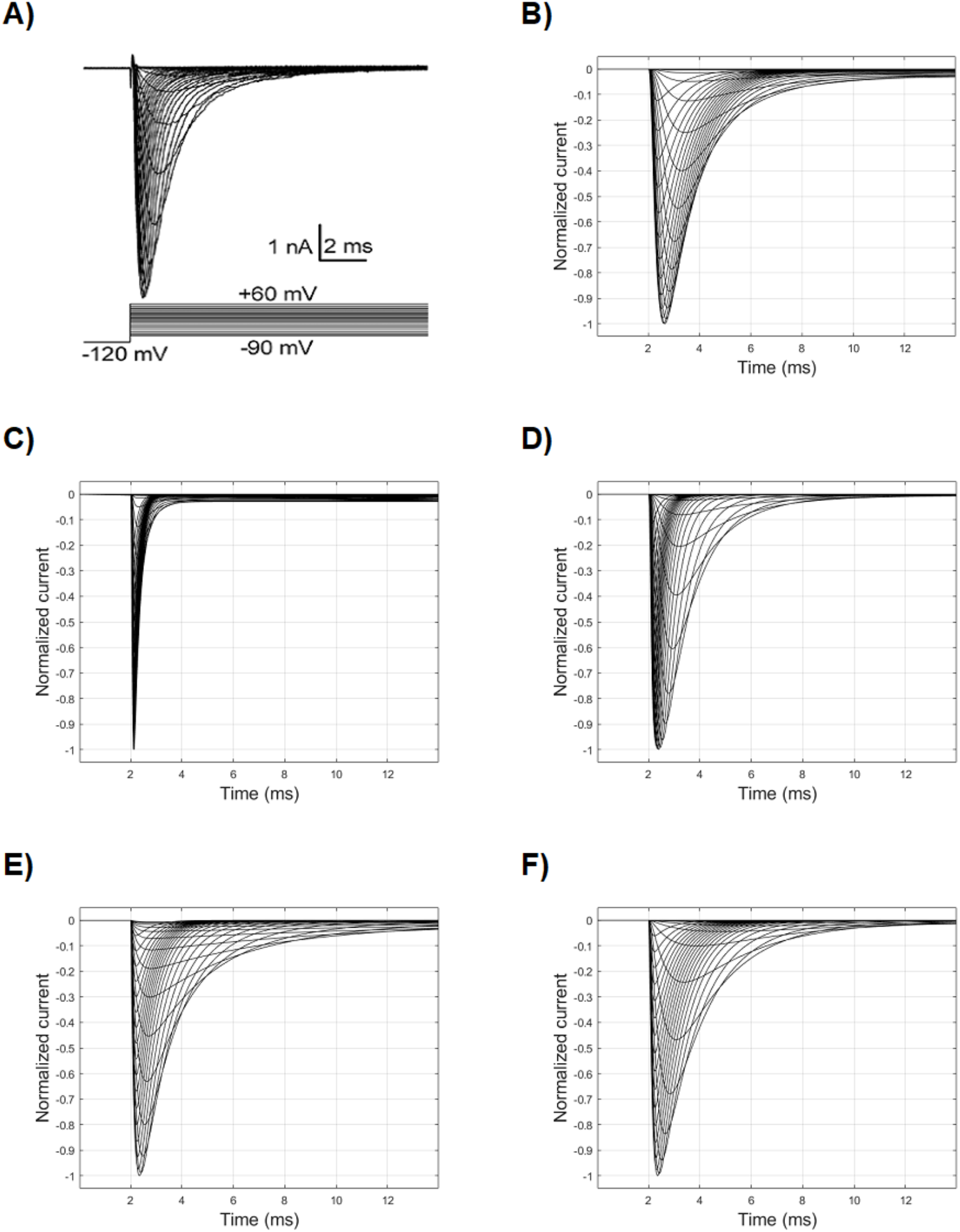
Real and simulated sodium current curves following the activation voltage clamp. A) Real curves from a previously published experimental study (Zhang et al, 2013), recorded at 25°C. B) Simulated curves obtained by the original HH model (Hodgkin and Huxley, 1952a), at 6°C (Table 2a). C) Simulated curves obtained by the original HH model, at 25°C. D) Simulated curves after modifying the original HH model (Table 2b). E) Simulated curves obtained by a kinetic model with 4 states: one closed, one open and two inactivated states. F) Simulated curves obtained by a kinetic model with 5 states.

**Table 2.**
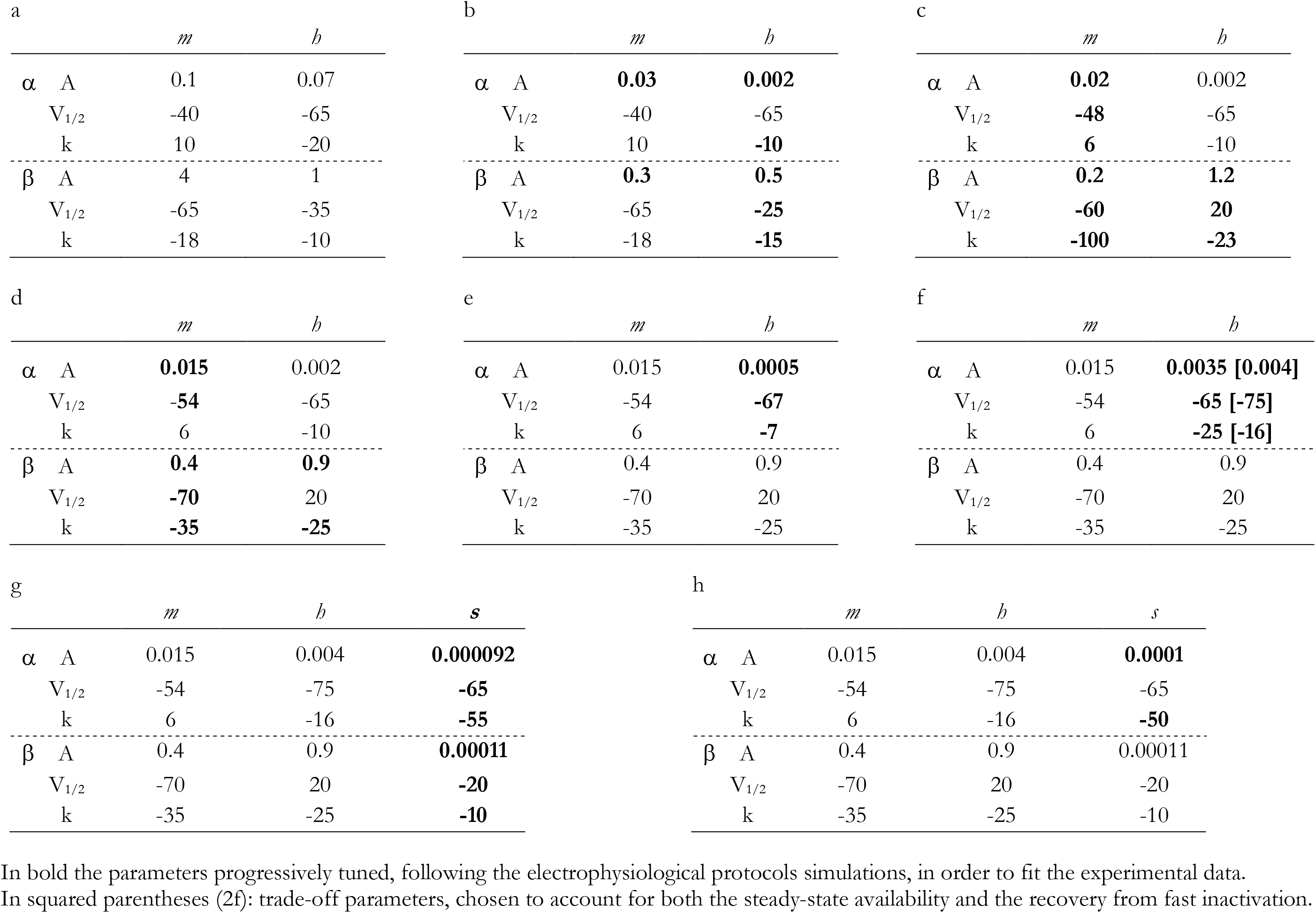
Parameters of the HH model.

In the process of empirical fitting, the reduction of both the time constants of activation (time to the peak of activation) and inactivation (decay from activation) was mainly achieved by decreasing the rate constants amplitudes, while the approximately correct sequence of activation and inactivation across the voltages clamp was achieved by shifting the *V_1/2_* of the backward rate constant of inactivation and by modifying the slopes of the rate constants of inactivation (Table 2b).

The voltage dependence of normalized conductance in Na_V_1.5 has flat or slightly decreasing values along its course for the most depolarized values (starting from 0 mV, approximatively; Fig 6A), while the corresponding values obtained with the just modified HH equations displayed an increasing course (Fig 6C).

**Figure 6.**
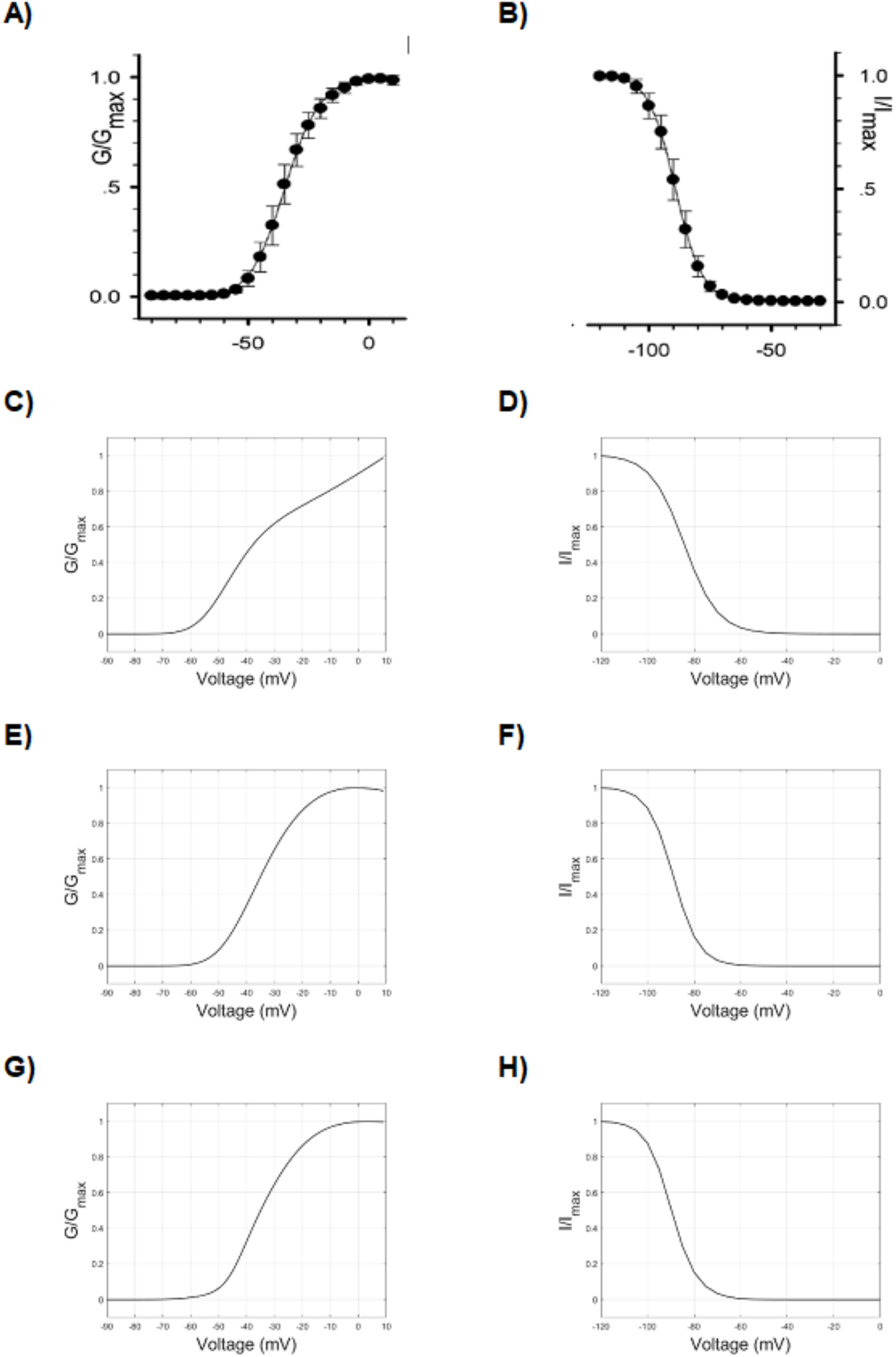
Voltage/Normalized conductance relationship during activation (left column) and Voltage/Normalized current relationship during fast inactivation (steady-state availability) (right column). A) and B) Experimental data (Zhang et al, 2013). C) Simulated values of Voltage/Normalized conductance relationship during activation obtained by the HH model before parameters tuning (Table 2b). D) Steady-state availability before parameters tuning (Table 2d) in the HH model. E) Simulated values of Voltage/Normalized conductance relationship during activation obtained by the HH model after parameters tuning (Table 2c). F) Steady-state availability after parameters tuning (Table 2e) in the HH model. G) and H) Simulated values obtained by the kinetic model.

The activation-voltage relationship is mainly sustained by the reciprocal interaction of voltage dependence of *α_m_* and *β_m_*, which gives rise to the voltage dependence of *m_∞_*, as depicted on Fig 1C.

However, the voltage dependence of the activation peak can also be modified by acting on the rate constants of inactivation, and the correct (experimental) course of activation can be better approximated by modifying the slope and increasing the amplitude of *β_h_*, which carry out a smaller value of *h* (that is, greater inactivation) for more depolarized voltages. In addition, after tuning the parameters of *α_m_* and *β_m_* (Table 2c), a modelled curve much more similar to the experimental one can be obtained (Fig 6E; *V_1/2_*=34.7 mV, *k*=−7.2).

The activation and inactivation time constants (Fig 7A-F) were similarly fitted in the above passages by tuning the same parameters. Both were derived from the equation (3c): the former was computed at different voltages on segments from the onset to the activation peak, while the latter on segments from the activation peak to the resting potential (decay from activation). It is worth mentioning here that the decay from activation is a first measure of inactivation.

**Figure 7.**
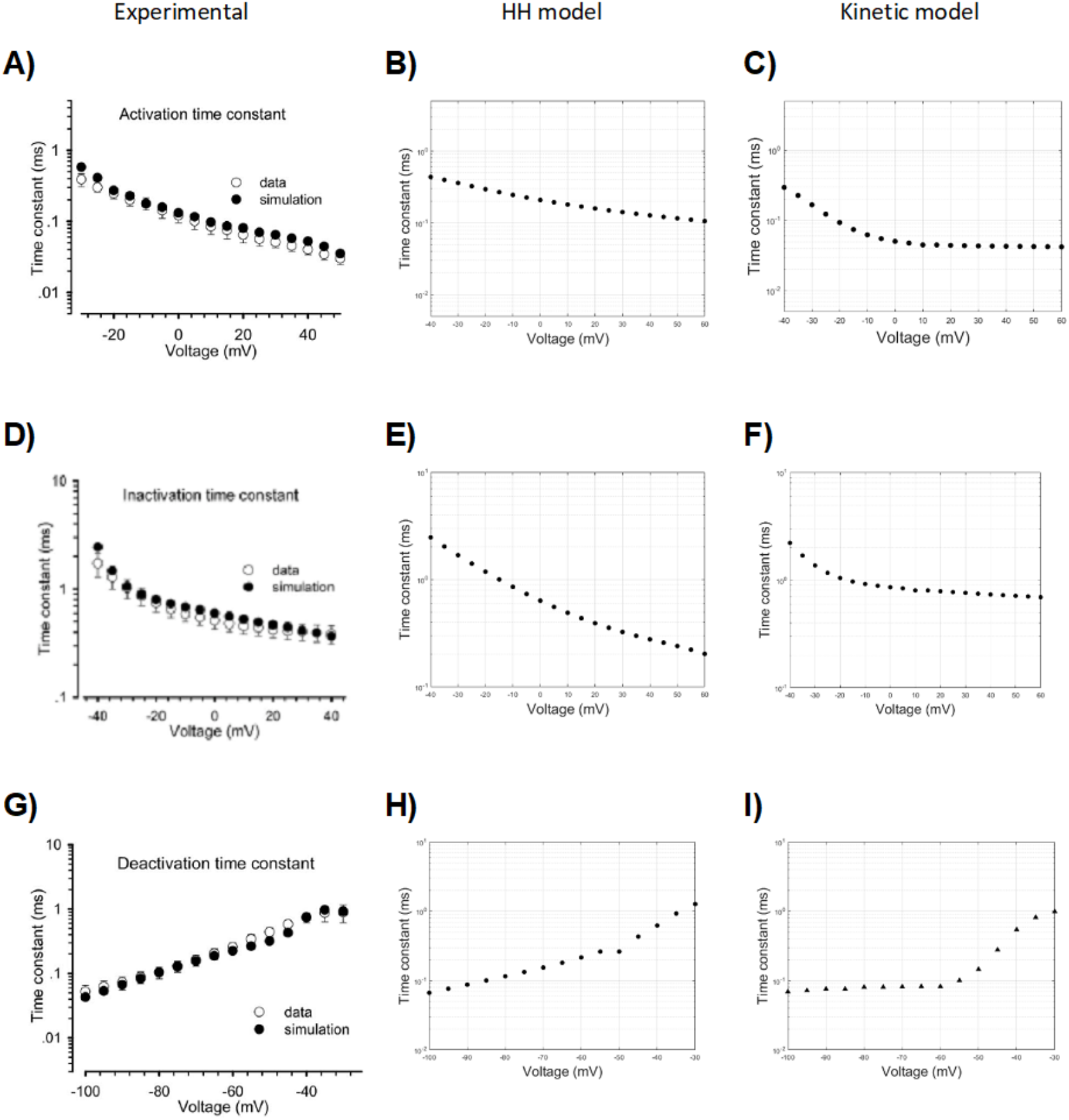
Time constants of activation (upper row), inactivation (middle row), and deactivation (bottom row) in function of voltage. Experimental values (Zhang et al, 2013) (left column), simulated data from a HH model (central column) and from a kinetic model (right column).

#### Kinetic model

The transition rates of the kinetic model were set in order to make all channels to be in the C1 state at polarized resting potential (Fig 3A-B; Table 3). By increasingly stepping the voltage towards more depolarized values, a progressively increasing fraction of channels moves to open conducting state (O1) through the second closed state (C2), due to increasing values of the transition rates between C1, C2 and O1. The forward and backward transition rates between C1 and C2, and between C2 and O1 determine the fraction of channels in O1 state and the time needed to reach this condition. However, since the O1 to I1 transition rate always has a non-zero value and the I1 to O1 transition rate is much smaller than the O1 to I1 transition rate, the fraction of channels in O1 state moves to I1 (that is, inactivates), with a velocity set by the O1 to I1 transition. The O1 to I1 transition, indeed, can be considered irreversible and the channel does not rest in O1, but moves to the inactivated state(s). With this kinetic model, the inactivated state follows the open state and is not independent from it, which is a behavior in agreement with real data (Bezanilla 2008).

**Table 3.** Parameters of the kinetic model.

| | $A_{hyper}$ | $V(1/2)_{hyper}$ | $k_{hyper}$ | $A_{dep}$ | $V(1/2)_{dep}$ | $k_{dep}$ |
| --- | --- | --- | --- | --- | --- | --- |
| *C1C2* | - | - | - | 8 | -16 | -9 |
| *C2C1* | 2 | -82 | 5 | 8 | -16 | -9 |
| *C2O1* | - | - | - | 8 | -26 | -9 |
| *O1C2* | 3 | -92 | 5 | 8 | -26 | -9 |
| *O1I1* | 8 | -50 | 4 | 6 | 10 | -100 |
| *I1O1* | 0.00001 | -20 | 10 | - | - | - |
| *I1C1* | 0.35 | -122 | 9 | - | - | - |
| *C1I1* | - | - | - | 0.04 | -78 | -10 |
| *I1I2* | - | - | - | 0.00018 | -60 | -5 |
| *I2I1* | 0.001825 | -88 | 31 | - | - | - |

The proposed kinetic model features a multi-step activation sequence, according to the experimental data (Patlak 1991), based on two closed states before the open state. Two closed states, indeed, are the minimum number of states able to replicate the activation curves and the voltage-dependence of normalized conductance (Fig 5F, Fig 6G, and Table 3). A too small, indeed, and unrealistic progression of the activation time constant results by adopting a single closed state (Fig 5E).

The open-to-inactivated transition (O1I1) contributes to the normalized conductance-voltage relationship and mainly sets the decay from activation.

### Deactivation

#### HH model

Tail currents (Fig 8A), evoked by a brisk repolarization after a very short depolarization pulse (before substantial channel inactivation), can be fitted to a monoexponential function (3d), whose time constant reduces progressively for more hyperpolarized test stimuli (Fig 7G). They are mainly modelled by the return of *m* at low values during repolarization. Following the previous adjustment of HH model parameters (Table 2c), the simulated curves exhibit too long time constants for more hyperpolarized test stimuli (Fig 8B). By varying the parameters of *β_m_* (Table 2d), which is responsible for low values of *m_∞_* at hyperpolarized values, the model is able to reach fairly similar time constants in the hyperpolarized range (Fig 8C). However, due to changes in the activation-voltage relationship induced by the *β_m_* variations, slight adaptations of *α_m_* and *β_h_* parameters (Table 2d) are needed to achieve again an acceptable fitting of the activation relationship (*V_1/2_*=34.4 mV, *k*=−7.2).

**Figure 8.**
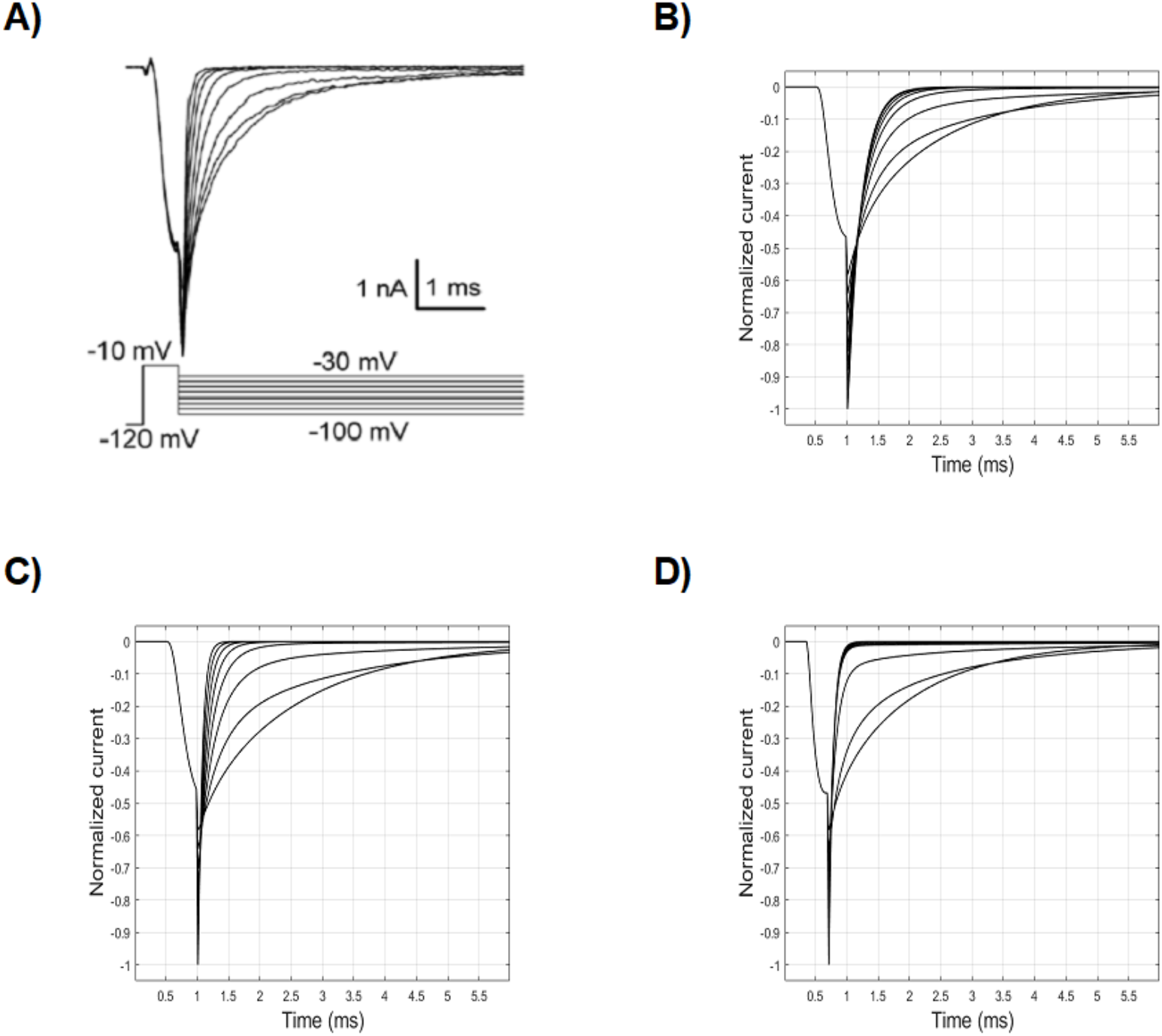
Current curves following the deactivation protocol. A) Experimental values (Zhang et al, 2013). B) Simulated values with the HH model before parameters tuning. C) Simulated values with the HH model after parameters tuning. D) Simulated values by means of the kinetic model.

#### Kinetic model

The transition from the open (O1) to the closed (C2) state (Fig 3D) is mainly responsible for the tail currents in the kinetic model. In particular, a fair approximation of experimental values (Fig 8D and 7I) can be reached by finely tuning the more hyperpolarized half of the transition curve. In that range, the prevalent fraction of closed channels is assured by the lower values of the corresponding C2O1 transition (Fig 3C).

### Fast inactivation

#### HH model

The voltage dependence of normalized current during fast inactivation (steady-state availability) represents the fraction of ionic channels not inactivated (i.e., available to activation), at different steady-state voltages. It is mainly given by the interplay between *α_h_* and *β_h_*, which carries out *h_∞_* values close to 1 at hyperpolarized voltages and close to 0 at depolarized voltages, as depicted in Fig 1B and Fig 1D. Experimental values of steady-state availability in Na_V_1.5 are depicted in Fig 6B. The HH parameters values set until now (Table 2d) reproduce an availability curve with fairly correct morphology (Fig 6D), although shifted towards too depolarized values (*V_1/2_*=84.1 mV), and too slanted (*k*=7.1). In order to shift the curve towards more hyperpolarized value (i.e. inactivation develops earlier, at more hyperpolarized voltages), it can be useful to reduce the amplitude of *α_h_*, while maintaining the value of *β_h_* unchanged, in order to not modify the reached good approximation of the activation curve. In addition, we can also try to modify the slope of *α_h_* to obtain a quicker transition to inactivation.

By adjusting the parameters of *α_h_* (Table 2e) it is possible to obtain a reasonable approximation (*V_1/2_*=−88.8 mV, *k*=5.5) to the HH experimental curve (Fig 6F). The parameters of the steady-state availability curve (that is, hemi-inactivation and slope) represent the second measure of the inactivation, distinct from the decay from activation. It is worth noting that *α_h_* mainly governs the steady-state availability (more hyperpolarized values), and *β_h_* is responsible for the activation relationship (for more depolarized voltages, as seen above), which also affects the decay from activation.

In addition, as understood by the steady-state availability curve, the value of *h* at hyperpolarized voltages (more negative than −100 mV) should be 1 (maximum, or quite so - that is, no inactivation). On the other hand, a quick inactivation develops at voltages around −90 mV, well before the start of activation (Fig 6A). As shown, both the phenomena can be addressed by finely tuning the interplay between *α_h_* and *β_h_*.

#### Kinetic model

In the electrophysiological protocol for studying the steady-state availability (Fig 4C), long enough conditioning stimuli of increasing voltage are able to set an increasing fraction of channels into an inactivated state, and the following test stimulus samples the fraction of channels available for activation (not inactivated). What is interesting and crucial here is that even conditioning stimuli below the activation threshold are able to promote the transition to an inactivated state. Therefore, at subthreshold depolarizations a fraction of channels transits from the closed state to the inactivated state, without passing through the open state. While the HH formalism does not explicitly model open, closed and inactivated states, the proposed kinetic model can specifically address this mechanism by tuning the C1 to I1 transition (C1I1, Fig 3D). The resulting curve for the kinetic model is displayed in Fig 6H, and the corresponding values of the C1I1 parameters are displayed on Table 3.

### Recovery from fast inactivation

#### HH model

The recovery from inactivation (or repriming) is sampled by a highly depolarizing conditioning pulse followed by a variable time interval of repolarization (from tenths to hundreds of milliseconds, in the case of Na_V_1.5), followed by a second depolarizing pulse, which tests the fraction of channels recovered from the inactivation (following increasing time intervals of repolarization) (Fig 4D).

During the first pulse of depolarization, *m* quickly increases (up to 0.8, in our simulation) (Fig 1C), while the *h* value, which starts at a value of 1 (polarized resting voltage, that is no inactivation), decreases to a value of 0 (Fig 1D) with a slower time constant compared to *m*. This provides the mechanism for the peak of sodium current. At the steady-state of activation-depolarization, which in Na_V_1.5 is reached within about 10 ms after the start of the conditioning pulse, *h* is equal to 0 (complete inactivation) and *m* is equal to 0.8 in our simulation, which produces no sodium current at all. The following repolarization, due to the higher value of *β_h_*, quickly sets the *m* value to 0 and, with a slower time course, the *h* value to 1 (absent inactivation).

By repolarizing the membrane with increasing intervals, the recovery from inactivation kinetics are sampled, which are mainly carried out by (the voltage dependence of) *α_h_*, provided that *β_h_* has values close to 0 at −120 mV (the repolarization voltage), and that the kinetic of *m* is much faster (Fig 1E).

Compared to the experimental recovery curve (Fig 9A, white circles), the one obtained by using the HH parameters set as above and displayed on Table 2e (Fig 9B) already shows an unrealistic substantial recovery after 0.1 ms of repolarization, and a much faster recovery time. By reducing the amplitude of *α_h_* (Table 2f), a more correct fitting can be obtained (Fig 9C). However, since the forward rate constant of inactivation *α_h_* also directly affects the fast inactivation curve, as shown above, the modified value does not provide a good fitting of the steady-state availability curve anymore (Fig 10C, to be compared to Fig 10A and Fig 10B).

**Figure 9.**
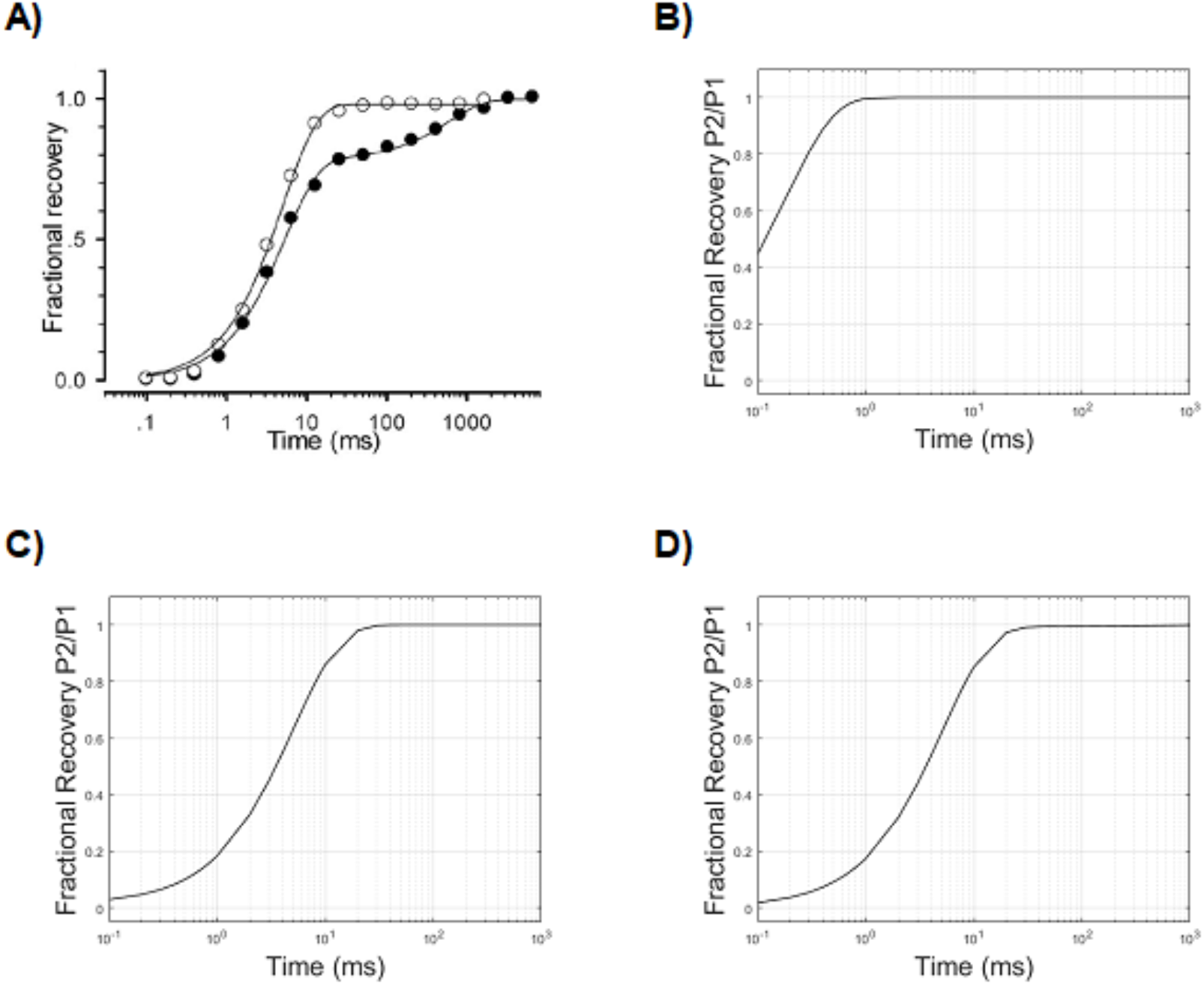
Recovery from fast inactivation, ratio P2/P1 in function of time intervals of repolarization (logarithmic axis). A) Experimental values (empty circle) (Zhang et al, 2013). B) Simulated values with the HH model before parameters tuning (Table 2e). C) Simulated values with the HH model after parameters tuning (Table 2f). D) Simulated values obtained by the kinetic model.

**Figure 10.**
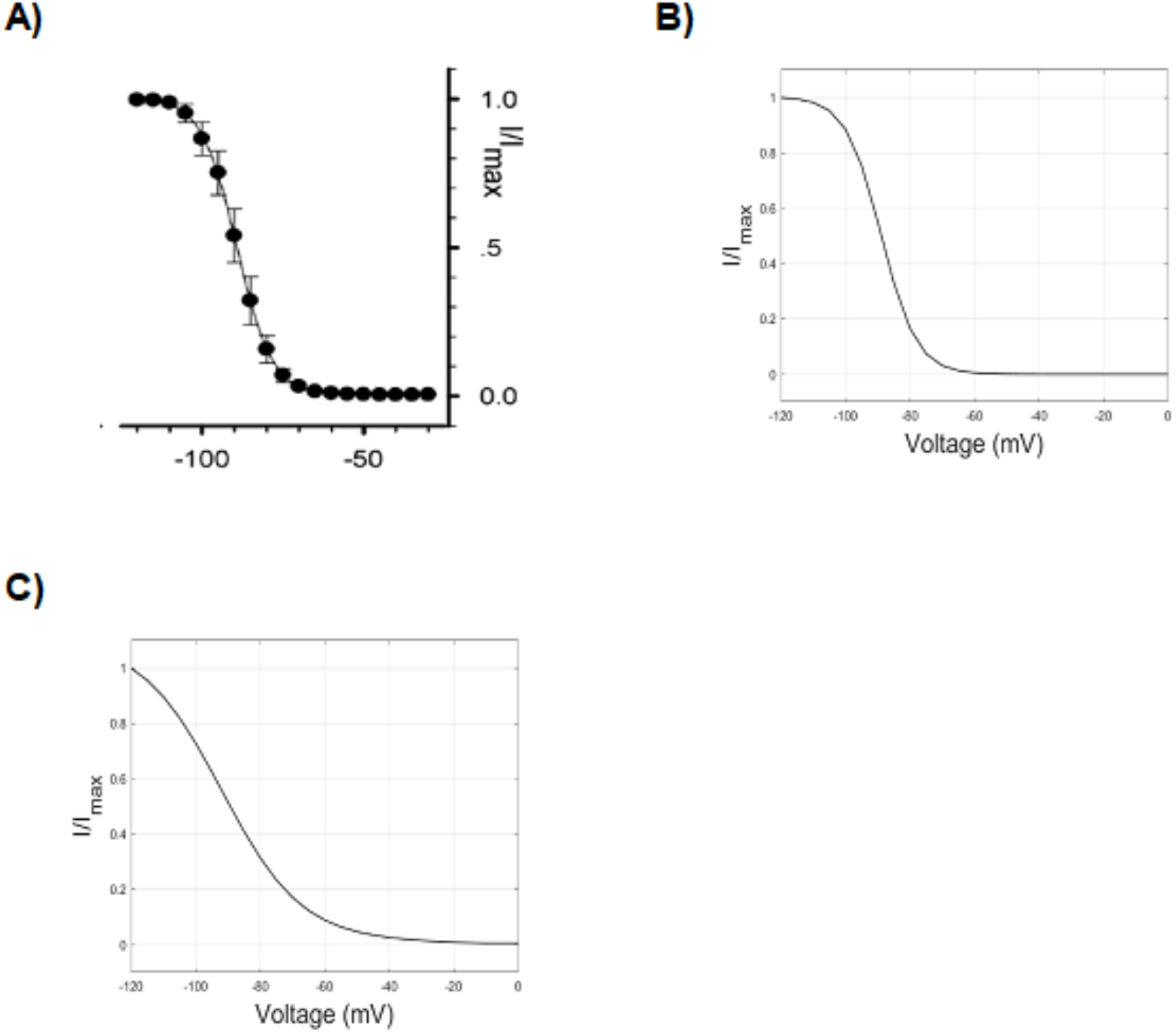
Effect of optimization of recovery from fast inactivation on steady-state availability in the HH model. A) Experimental values of Voltage/Normalized current relationship following steady-state availability protocol (Zhang et al, 2013). B) Simulated values of Voltage/Normalized current relationship following steady-state availability with the HH model before tuning parameters (namely *α_h_*) in order to optimize the recovery from inactivation (Table 2e). C) The same relationship after parameters tuning (Table 2f).

Thus, the voltage dependence of *α_h_* in HH models affects both the recovery from inactivation and the steady-state availability. This results in a conflicting limitation for the parameter optimization of the model. In other words, the detailed reproduction of one electrophysiological feature (the steady-state availability) affects the accuracy of the other (the recovery from inactivation). In Table 1, an example of this conflicting behaviour of the model is provided: the parameters of *α_h_* have been set to preferably reproduce in detail the repriming (a), or the steady-state availability (b); alternatively, a trade-off can be made, by choosing parameters which carry out the closest, yet not detailed, approximation for both the electrophysiological features (c).

In addition, even when the HH model is set to reproduce the time constant of repriming, it fails when different voltages of repolarization are chosen (Table 1). In order to fix this incorrect behaviour, one should also modify *β_h_*, which has relevant effects (i.e., it affects the time constant of recovery) in the hyperpolarized range (around −100 mV) when the value of *α_h_* is small. However, each variation of *β_h_*, as mentioned above, also changes the voltage dependence of activation, which, in this case, results not amendable by further modifying *α_m_* and *β_m_* (data not shown).

Finally, it is worth mentioning that the recovery from inactivation provides a third measure of the fast inactivation.

#### Kinetic model

Recovery from inactivation can be easily set in detail by tuning the transition from the inactivated state I1 to the closed state C1 (I1C1, Fig 3D), a way which is currently considered much more alike to the real biophysics of the channel (Fig 9D; Table 3). In addition, no conflict arises with the electrophysiological behavior evoked by the steady-state availability protocol, which is mainly governed by the C1I1 transition, as shown above.

### Development of slow inactivation

#### HH model

In addition to fast inactivation, Sodium channels also exhibit a slower inactivation, which was unrecognized when the HH model was developed.

The development of the slow inactivation can be sampled by the protocol depicted on Fig 4E.

To reproduce the slow inactivation kinetics by means of the HH formalism, an additional third particle (or gate) has to be considered in the conductance equation (refer to equation 4), which is usually named s and carries rate constants formally similar to those of *h*,

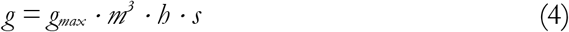

In this case, the entry into the slow inactivation is mainly modelled by tuning the voltage-dependence of the *α_s_* rate constant (which, similarly to *h*, controls the transition from 1 to 0 of *s*). Also, it should be considered that, for the depolarization voltage (P2) used in the protocol (that is, −20 mV), even *β_s_* must be tuned (that is, not 0 value), in order to obtain the correct proportion of non-slowly-inactivating channels even at longer durations of P1.

Therefore, after adding the *s* factor to the HH model and tuning its parameters (Table 2g), a fairly good approximation of the development of slow inactivation can be reached (Fig 11C). Note that, for this simulation, an intermediate setting of *α_h_* has been adopted to provide a trade-off between the best fittings of steady-state availability and recovery from fast inactivation.

**Figure 11.**
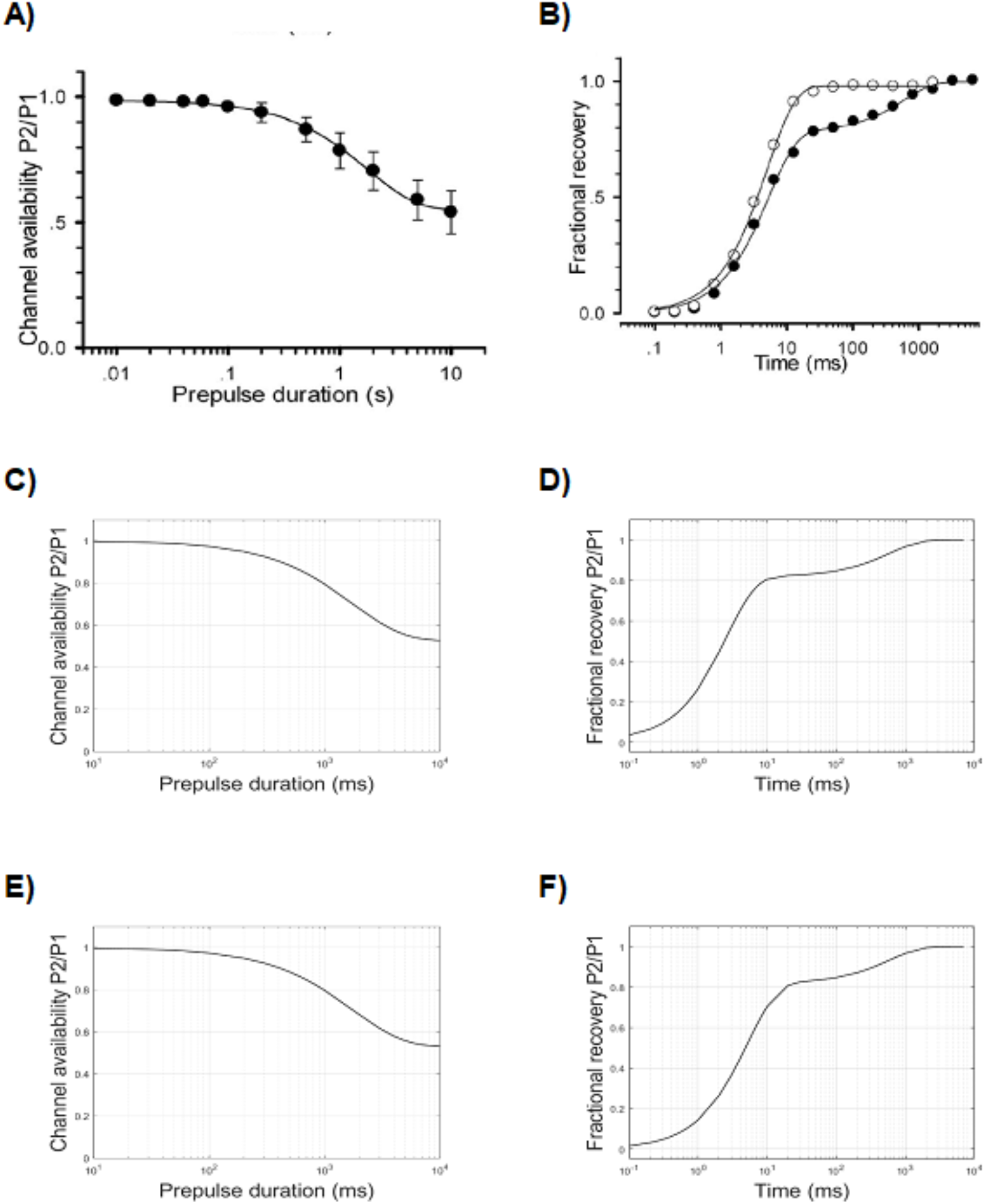
Development of slow inactivation (left column) and recovery from slow inactivation (right column). Abscissae in logarithmic scale of time intervals. A) Development of slow inactivation, and B) recovery from slow inactivation (solid circle) in the experimental setting (Zhang et al, 2013). C) Development of slow inactivation (Table 2g), and D) recovery from slow inactivation (Table 2h) in the HH model. E) Development of slow inactivation, and F) recovery from slow inactivation in the kinetic model.

#### Kinetic model

The development of slow inactivation in the kinetic model is easily approximated by tuning the parameters of I1 to I2 transition (Table 3, Fig 3E). Analogously to the HH model, to reach the correct proportion of not-slowly-inactivating fraction of the channels, even the I2 to I1 transition should be tuned, in a way that at −20 mV (the conditioning voltage) its value is different from 0.

The obtained modelled development of slow inactivation is depicted in Fig 11E.

### Recovery from slow inactivation

#### HH model

Similarly to the fast inactivation, also the recovery from slow inactivation is sampled by means of a double pulse protocol (Fig 4E), in which the first depolarizing conditioning pulse (P1) is followed by a repolarizing time interval of increasing duration (Δt, from 0.1 s to 10 s), followed by a short probing depolarization (P2). The difference with the fast inactivation protocol is that P1 is much longer (1000 ms, compared to 100 ms).

By adopting this protocol, a recovery time course develops with two time constants of recovery, the first and faster one for the kinetics of fast inactivation, and a slower one for the slow inactivation repriming. By tuning the slow inactivation (*s*) rate constants (Table 2h), a good agreement with the recovery time constant of slow inactivation can be obtained (Table 1 and Fig 11D). Note that the time constant of recovery from fast inactivation is not correct because we adopted, for these simulations, an intermediate setting of *α_h_* (Table 2h).

#### Kinetic model

The recovery from slow activation can be easily reproduced in detail by finely tuning the I2I1 transition (Fig 3E). The low values of the amplitude parameter of I1I2 and I2I1 stand for the slow processes of entry into and recovery from this kind of inactivation. The resulting curve of repriming of the slow inactivation is depicted in Fig 11F).

### Channel models implementation in a standard neuron model

This subsection is only intended as a not exhaustive proof of concept of the feasibility and suitability of the proposed channel model to be implemented in neuron computational models, mainly performed in order to test their computational load. To this aim, we exploited a reduced and simple neuron computational model, previously developed by Dodge and Cooley (Dodge and Cooley 1973). Due to the limited purpose of this simulation, no in-depth exploration of the implementation has been performed, and also a series of approximations, yet not realistic, have been adopted (e.g., the implementation of the heart sodium channel in a spinal motoneuron model, even though Na_V_1.5 has been also detected in the central nervous system (Wu et al. 2002)). The model developed by Dodge and Cooley is a reduced model of spinal motoneuron equipped with sodium, potassium and leakage conductances and comprises a soma, an equivalent dendrite, an initial segment, a myelinated segment, a node, and an extra unmyelinated segment. In order to sample the suitability and versatility of our models, we simply substituted the original HH sodium conductance mechanism with either the modified HH model or the kinetic one, tuned with the parameters specified on tables 2h and 3, respectively.

Apart from the substitution of the original sodium conductance with our Na_V_1.5 models, we only set the resting potential (and in turn the leakage equilibrium potential) to −100 mV, instead of −80 mV, to take into account the different kinetics of Na_V_1.5 inactivation. By maintaining unaltered all other biophysical and structural parameters, the neurons carrying the HH and kinetic channel models, when stimulated by a virtual electrode located into the soma, were able to fire spikes, although the action potentials showed a wider duration compared to the Dodge and Cooley model (see Supplemental files, Fig S1A-C), and an increased threshold of the stimulus (~150 nA versus ~65 nA). In order to compare the computational load between the models, a 3000 ms long simulation was performed, in which a train of electrical stimuli (2800 ms long, with an amplitude of 160 nA and a frequency of 25 Hz) was delivered to the soma. In this simulation, a time step of integration of 0.001 ms was chosen, and ten runs were performed for each channel model.

Despite our expectations, the neuron model equipped with the HH channel model completed the simulations only about 2 s before the one equipped with the kinetic channel model (34.19 ± 0.34 s versus 36.38 ± 1.43 s). Because of the minor computational load of the HH formalism we would have expected a greater difference. The original Dodge and Cooley model, equipped with channels built according to the HH formalism, completed the runs in 31.50 ± 0.14 s (Table 4).

**Table 4.**
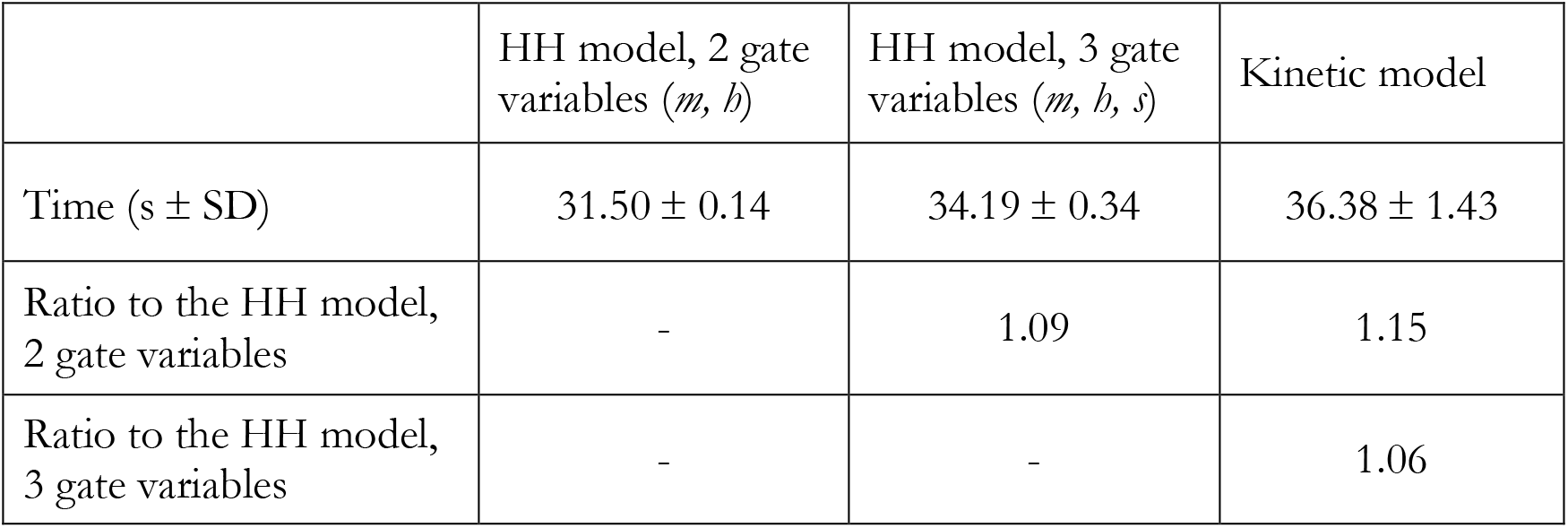
Running time values.

## Discussion

Our study recognizes some critical limitations of the HH formalism in modelling the known complexity of Na_V_1.5 macroscopic currents, and shows how a simplified kinetic model is better suited to approximate in detail these currents. It also provides a practical guide to a procedural optimization of simplified kinetic models, which are shown to exhibit a computational load comparable to that of the HH model.

The need to develop models of ionic channels able to reproduce in detail more recent electrophysiological data has been for long recognized (Cannon and D’Alessandro 2006; Destexhe and Huguenard 2010). On the other hand, both theoretical (Patlak 1991; Kuo and Bean 1994; Bezanilla 2008) and practical (Strassberg and Defelice 1993; Meunier and Segev 2002) limitations of HH models have been already reported. Among these, the HH model hypotheses of independent gating particles, the calculated time constant of deactivation, the behaviour of the gating current, have been proven inaccurate (Sterrat et al. 2011).

In addition, here we demonstrate a critical practical limitation of the HH model which, to the best of our knowledge, has not been previously put explicitly forward. The steady-state inactivation and the recovery from inactivation have to be set in HH models by tuning an identical and unique parameter, the voltage dependence of *α_h_*. Thus, the detailed reproduction of the two electrophysiological behaviours results in a conflicting optimization limit.

In one of their seminal papers (Hodgkin and Huxley 1952b), Hodgkin and Huxley devised three electrophysiological protocols to specifically study the behaviour of the (fast) inactivation of the voltage-gated sodium channel of *Loligo*. The protocols (Fig 4C-E), with few modifications, would have been later known as steady-state availability, recovery from inactivation (or repriming) and development of inactivation, and, as for the entire Hodgkin and Huxley research, they set the framework for all following studies on the subject. In that paper, poor resolving power of the material (*Loligo* sodium channel has time constants of entry into inactivation - from closed states - close to the time constants of decay from activation) likely prevented the Authors to recognize that entry into inactivation can proceed along different mechanisms (namely from closed or open states). Consequently, they suggested only one time constant (named *τ_h_*) to describe the inactivation processes (Hodgkin and Huxley 1952a, b). Thus, in HH models *β_h_* (Fig 1B) usually sets the decay from activation (due to its high value in depolarizing range) and *α_h_*, which properly establishes the recovery from inactivation, must be also used for providing the inactivation in a less depolarized range.

With the proposed kinetic model, on the contrary, each variable can be set independently: the decay from the activation by the O1I1 parameters, the recovery from activation by the I1C1 parameters, the steady-state availability by the C1I1 parameters.

### The phenomenological modelling

We recently described how a single simplified kinetic model is able to reproduce in detail the macroscopic currents of all human voltage-gated sodium channels, Na_V_1.1 to Na_V_1.9 (Balbi et al. 2017). Apart from the adherence to different experimental data, the proposed phenomenological model was also constrained by the parsimony of states and transitions, in order to build the simplest framework with minimal computational load and to provide a kinetic model suitable for the implementation in large conductance-based neural networks.

The present study only deals with the macroscopic currents, and the proposed kinetic model is not intended to fit the data from biophysical studies of the channel, where single channel recordings are exploited to derive the subtle microscopic conformational changes of the pore-forming protein (e.g., Börjesson and Elinder 2008), also taking into account the gating currents. In the present study the single states do not correspond to physical molecular states of the protein, and they should be rather considered as aggregates of molecular configurations operationally grouped into a set of distinct states separated by large energy barriers (Hille 1992). For example, while a series of four (or more) closed states are commonly hypothesized (usually in adherence with the tetrameric structure of the proteic channel) before an open state can develop following a depolarizing step, our model collapses them all in only two. Two closed states, indeed, are necessary and sufficient to deterministically reproduce the phenomenological behaviour of the channel.

### Theoretical and practical advantages of the kinetic model

At variance with HH formalism, kinetic models, yet simplified, try to not contradict what is known about the biophysics of the channel, as understood by functional and structural studies. For example, they can account for the direct transition from closed to inactivated states (Patlak 1991), for the inactivation passing through the closed states before reactivation (Kuo and Bean 1994), for the multistep process of activation (Patlak 1991), for the dependence of inactivation from activation (Bezanilla 2008).

In addition, Markov-type kinetic models appear to be more adaptable and prone to incorporate novel or further channel features with respect to HH models. Indeed, as a general computational tool, they have been already exploited to simulate different neural phenomena, like ligand-gated channels or second messenger-activated channels, and they have been considered as a more general framework in the larger context of biochemical signal transduction (Destexhe et al. 1994).

The results from the channels implementation in a neuron model show that the runtime of the kinetic model is only about 5% higher than that of the HH model, which seems a completely repayable drawback, considering the advantages that the kinetic model provides in terms of efficacy, level of approximation to the experimental electrophysiological data, and adherence to the biophysics of the ionic channels.

### Predictions and suggestions from the kinetic model

Modelling studies can suggest slight variations of the canonical electrophysiological protocols, in order to clarify some less known features of the channels. For example, by adopting different levels of repolarization voltages during recovery from inactivation protocols (Zhang et al. 2013), a fractional recovery can be recognized (Table 1). Analogously, even the steady-state availability protocol could take advantage by adopting different levels of conditioning steady-state voltages.

As regarding the evaluation of time constants of activation and inactivation, we fitted the curve obtained by the activation protocol with the equation (3c), which involves a third power exponential to fit the activation segment of the curve, and a simple exponential for the inactivation (decay) segment. In the study carrying the experimental values we refer to (Zhang et al. 2013), a slightly different fitting procedure was adopted. Two separate fits for the activation and inactivation segments were performed, and both of them were fitted to a simple exponential, after (presumably) manually splitting the two segments of the curve.

Although this different fitting approach brought results slightly divergent (Fig 7) from those by our experimental reference (Zhang et al. 2013), we chose it because: a) it is the original procedure from the work by Hodgkin and Huxley (Hodgkin and Huxley 1952a), b) it provides the best fitting (minimum fitting error), c) it is free from errors derived by manually splitting the curves.

In addition, in the present study a measure of the fitting error has been always reported. This is unfortunately an uncommon practice in ion-channels modelling, which, instead, could greatly improve the reliability of the comparison among models and their refinement.

### Limitations and future studies

The major limitation of the present modelling study comes from the lack of raw electrophysiological data. It would have been better to optimize the models based on raw clamping currents data instead of indirect relationships, such as normalized conductance-voltage relationship following activation voltage clamping, normalized current-voltage relationship following fast inactivation protocol, etc. Moreover, the possibility to compare the simulated data directly to the raw data would provide a more explicit and reliable model testing.

The availability of raw electrophysiological data in open access web repository with standard and shared format could overcome this limitation, common to other modelling studies, and would greatly improve the quality of ionic channels modelling studies.

Modelling represents the challenging, continuous effort to approximate some physical phenomena, given some established or hypothetical mathematical description of it. In this sense, it is a matter of progressive approximation and continuous updating, according to the deepening knowledge of the examined phenomena. The present study shows how the adoption of kinetic models, along with the effort to limit their complexity (the ‘Occam’s razor’ principle), could also advantage the biologically inspired modelling of other ion channels, both voltage- and ligand-gated.

In following studies, further comparisons between HH and kinetic models could be undertaken by exploiting more detailed cell models and neural networks. Previous studies (e.g., Maurice et al. 2004) already developed kinetic models of ionic channels in order to fit experimental data not accounted for by HH models in detailed neuron cell models. Due to the limitations of HH formalism in modelling the inactivation kinetics, post-spike refractory times could show the most striking differences with kinetic models.

## Information Sharing Statement

The source code developed in NEURON 7.6 simulation environment, comprehensive of channel models in NMODL and all virtual experimental procedures, is available as a ModelDB (McDougal et al. 2017) entry (access number: 257747) (http://modeldb.yale.edu/257747).

## Aknowledgements

We thank prof Jeanette Hellgren Kotaleski and prof Paolo Massobrio for their comments and suggestions on a preliminary draft of the study.

---

**Figure S1.**
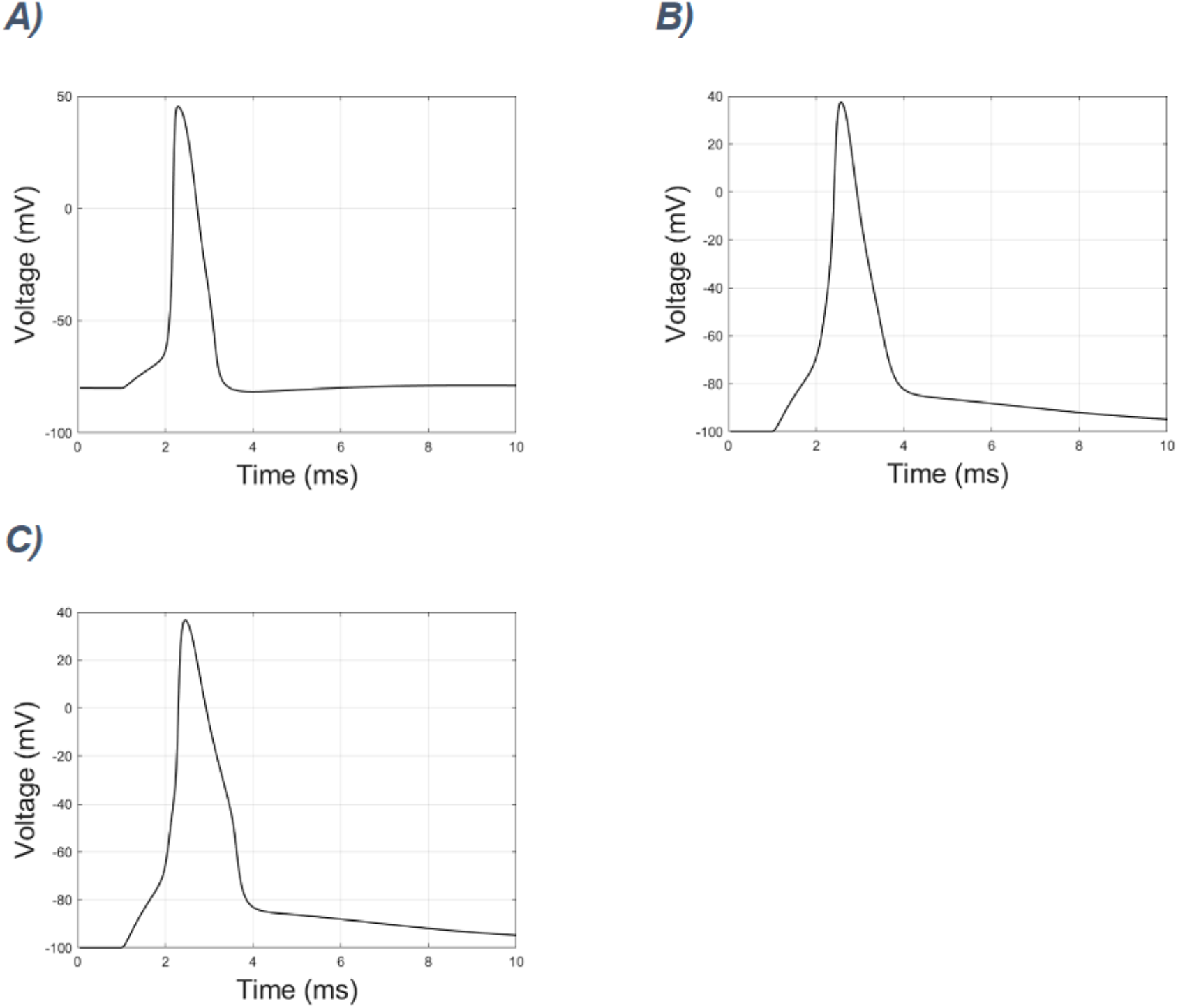
Action potential in a reduced spinal motoneuron (Dodge and Cooley, 1973) following a depolarizing somatic stimulus. A) Voltage displacement at the soma of the original cell model. B) Voltage displacement at the same location after substituting the original model of sodium channel with the Na_V_1.5 model built according to the HH formalism, or C) with the Na_V_1.5 kinetic model.

